# A One-Step Chemoselective Strategy for Hydroxymethylcytosine Sequencing in DNA and RNA

**DOI:** 10.64898/2026.09.14.751533

**Authors:** Jiahao Li, Pei-Hong Zhang, Yanchu Arvin Wang, Yuhao Zhong, Yiding Wang, Ruitu Lyu, Chenyou Zhu, Qing Dai, Chuan He

**Author notes:** These authors contributed equally (Jiahao Li, Pei-Hong Zhang). **Corresponding Author** Chuan He – Department of Chemistry, Department of Biochemistry and Molecular Biology, Institute for Biophysical Dynamics, The University of Chicago, Chicago, IL, USA; Howard Hughes Medical Institute, The University of Chicago, Chicago, IL, USA.; Qing Dai – Department of Chemistry, Department of Biochemistry and Molecular Biology, Institute for Biophysical Dynamics, The University of Chicago, Chicago, IL, USA; Howard Hughes Medical Institute, The University of Chicago, Chicago, IL, USA.

## Abstract

5-Hydroxymethylcytosine (5hmC) is a relatively stable chemical mark formed by oxidation of 5-methylcytosine in DNA, while its RNA counterparts, 5-hydroxymethylcytidine (hm^5^C) and 2”-*O*-methyl-5-hydroxymethylcytidine (hm^5^C*m*), have recently emerged as potential regulators of post-transcriptional processes. Existing 5hmC enrichment methods rely on antibodies or enzymatic *β*-glucosyltransferase labeling, are largely DNA-specific, and are not readily applicable to RNA. Here we report bisulfite-assisted labeling by thiol sequencing (BALT-seq), a one-step chemoselective strategy that installs a biotin-thioether handle directly onto hydroxymethylcytosine in DNA or RNA under mild acidic bisulfite conditions, without using enzymes. Mechanistic studies show that thiol capture of the bisulfite-activated 5hmC intermediate competes with canonical cytosine-5-methylenesulfonate formation, yielding a stable adduct that DNA polymerases bypass efficiently and that resists APOBEC-mediated deamination. In mouse embryonic stem cells, BALT-seq produces genome-wide 5hmC profiles highly concordant with 5hmC-Seal, TAB-seq, and oxBS-seq, recovers expected gene-body and chromatin-state enrichment, and extends to low-input cell-free DNA. On RNA, BALT-seq enriches hm^5^C/hm^5^C*m*-containing transcripts more effectively than antibody-based hMeRIP and reveals structured modification landscapes across tRNA species and LINE/LTR repeat families. BALT-seq thus establishes a unified, enzyme-free chemical platform for hydroxymethylcytosine profiling across DNA, cell-free DNA, and RNA.

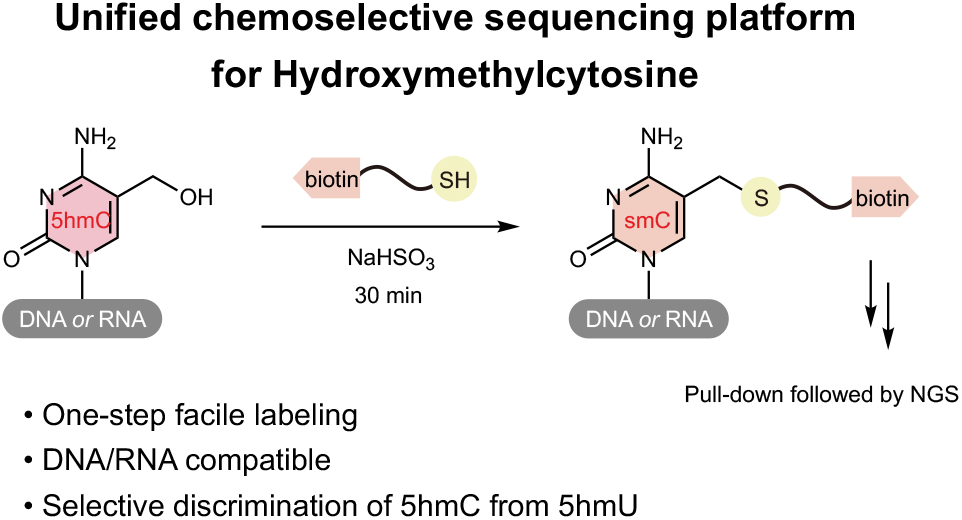

## INTRODUCTION

Cytosine modifications provide a chemically diverse layer of information on top of the genetic sequence. In mammalian DNA, 5-methylcytosine (5mC) is installed by DNA methyl-transferases and is dynamically remodeled through oxidation by the ten-eleven translocation (TET) family of Fe(II)/*α*-ketoglutarate-dependent dioxygenases. The first oxidation product, 5-hydroxymethylcytosine (5hmC), was recognized as a mammalian DNA base concurrent with the discovery of TET1 as the enzyme responsible for converting 5mC to 5hmC.^1,2^ Subsequent studies established that 5hmC can be further oxidized to 5-formylcytosine (5fC) and 5-carboxylcytosine (5caC), which can be processed by thymine-DNA glycosylase and base excision repair to complete active DNA demethylation (**Figure 1A**).^3^ 5hmC, however, is not simply a transient intermediate. It persists at measurable levels in many mammalian tissues, is especially abundant in the nervous system, and displays characteristic enrichment in gene bodies and regulatory elements.^4,5^

**Figure 1.**
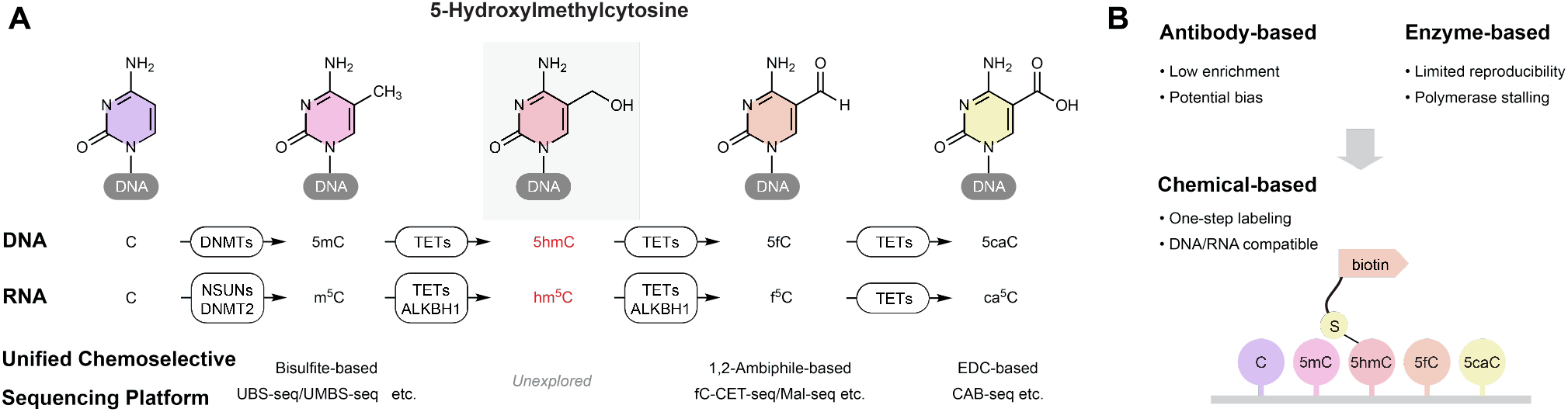
Unified chemoselective 5-hydroxymethylcytosine sequencing platform. (A) Cytosine modification pathways in DNA and RNA and enzyme-mediated oxidation pathways in epigenetic and epitranscriptomic regulation. (B) Conceptual design of bisulfite-assisted labeling by thiol (BALT), enabling sequencing-based detection of 5-hydroxymethylcytosine-containing DNA and RNA through direct covalent labeling.

The biological importance of 5hmC has created a continuing need for robust mapping technologies. 5hmC depletion is observed in multiple hematologic and solid tumors, and 5hmC patterns in genomic DNA and circulating cell-free DNA (cfDNA) have been used to infer disease state and tissue-of-origin information.^6–11^ Because 5hmC is often present at much lower abundance than unmodified cytosine and 5mC, enrichment-based methods are attractive for practical sequencing applications. Such methods reduce the sequencing depth needed to detect informative regions, can accommodate lower input amounts, and are compatible with high-throughput sample processing. Nevertheless, the quality of biological inference depends strongly on the chemistry used to distinguish 5hmC from related cytosine derivatives.

The most widely used enrichment strategies can be grouped into antibody-based and enzyme-based approaches. Direct immuno-precipitation with anti-5hmC antibodies is experimentally simple but is susceptible to epitope-density requirements, sequence or structural bias, and variable recovery.^12^ 5hmC-Seal uses T4 *β*-glucosyltransferase (T4 *β*GT) to transfer an engineered azido-glucose to 5hmC, followed by click chemistry and biotin pull-down.^13,14^ This approach provided sensitive genome-wide maps and has been widely adopted. However, it still requires an enzyme, a modified UDP-glucose donor, a click reaction, and multiple purification steps. The appended glucosyl-triazole-biotin group can also be bulky for downstream polymerase readout. Furthermore, *β*-glucosyltransferase chemistry is not compatible with RNA hm^5^C and hm^5^C*m*, and its specificity is not strictly limited to 5hmC in Watson-Crick-paired DNA; 5-hydroxymethyluracil-containing mismatches have been reported as potential substrates, creating a possible source of false-positive enrichment.^15^

Hydroxymethylcytosine modification is not restricted to DNA. In RNA, 5-methylcytidine (m^5^C) can also undergo oxidative processing to 5-hydroxymethylcytidine (hm^5^C), with emerging evidence suggesting dynamic regulation by TET enzymes and ALKBH1 (**Figure 1A**).^16,17^ A second hydroxylated derivative, 2”-*O*-methyl-5-hydroxymethylcytidine (hm^5^C*m*), has also been detected in mammalian RNA.^18,19^ Recent work has further connected RNA m^5^C oxidation to chromatin regulation and leukemia-relevant transcriptional states, emphasizing that oxidized RNA cytosines may have regulatory roles beyond canonical tRNA modification biology.^20^ Yet the analytical methods available for RNA hm^5^C/hm^5^C*m* lag behind those for DNA 5hmC. A major challenge is that RNA hm^5^C/hm^5^C*m* is not a substrate for *β*-glucosyltransferase, the key enzyme in established DNA 5hmC profiling strategies, rendering these approaches inapplicable to RNA. Moreover, antibody-based hMeRIP-seq enables transcriptome-wide enrichment of hm^5^C-containing transcripts.^21^ However, its dependence on antibody recognition and lack of chemical or base-resolution specificity limit confident definition of novel hm^5^C/hm^5^C*m* landscapes. A direct chemical enrichment method that operates on both DNA and RNA would therefore provide a useful bridge between epigenomic and epi-transcriptomic studies.

Past studies have established chemoselective strategies for several oxidized cytosine derivatives in both DNA and RNA by directly exploiting their intrinsic chemical reactivity. Bisulfite-based methods, including ultrafast bisulfite sequencing (UBS-seq)^22^ and ultra-mild bisulfite sequencing (UMBS-seq)^23^, have enabled base-resolution mapping of 5-methylcytosine through similar chemical conversion. 5-formylcytosine has been interrogated using selective 1,2-ambiphile chemistry, exemplified by fC-CET-seq^24^ and Mal-Seq^25^, whereas 5-carboxylcytosine has been mapped using carbodiimide-mediated activation strategies by EDC such as CAB-seq^26^. Hydroxymethylcytosine, however, has remained the missing piece in the development of a chemoselective sequencing platform for cytosine modifications across nucleic acids.

To address the challenge, we designed bisulfite-assisted labeling by thiol (BALT) to repurpose bisulfite chemistry for selective functionalization of hydroxymethylated cytosines that include DNA 5hmC and RNA hm^5^C and hm^5^C*m* (**Figure 1B**). We reasoned that if the bisulfite-activated 5hmC intermediate could be intercepted by an exogenous thiol bearing biotin, the reaction would generate a direct affinity tag in one step. This strategy would avoid antibodies and enzymes while retaining the high sensitivity of affinity enrichment.

## RESULTS AND DISCUSSION

The chemical design of BALT was guided by an unusual behavior of 5hmC under bisulfite treatment. Classical bisulfite sequencing treats nucleic acids under strongly converting conditions so that cytosine is deaminated to uracil, whereas 5mC is relatively resistant. 5hmC reacts with bisulfite in a different way, forming cytosine-5-methylenesulfonate (CMS), a property previously exploited by anti-CMS mapping.^27,28^ This observation suggested that the C5 substituent of 5hmC could be converted into a chemically addressable electrophilic intermediate, therefore thiols can be introduced as external nucleophiles (**Figure 2A**).^29,30^ Based on these observations, we asked whether thiol capture could outcompete bisulfite trapping to yield a stable thioether product and whether reaction condition optimization can lead to fast and almost quantitative conversion to thioether product as an effective way for selective 5hmC chemical labeling (**Figure 2A**).

**Figure 2.**
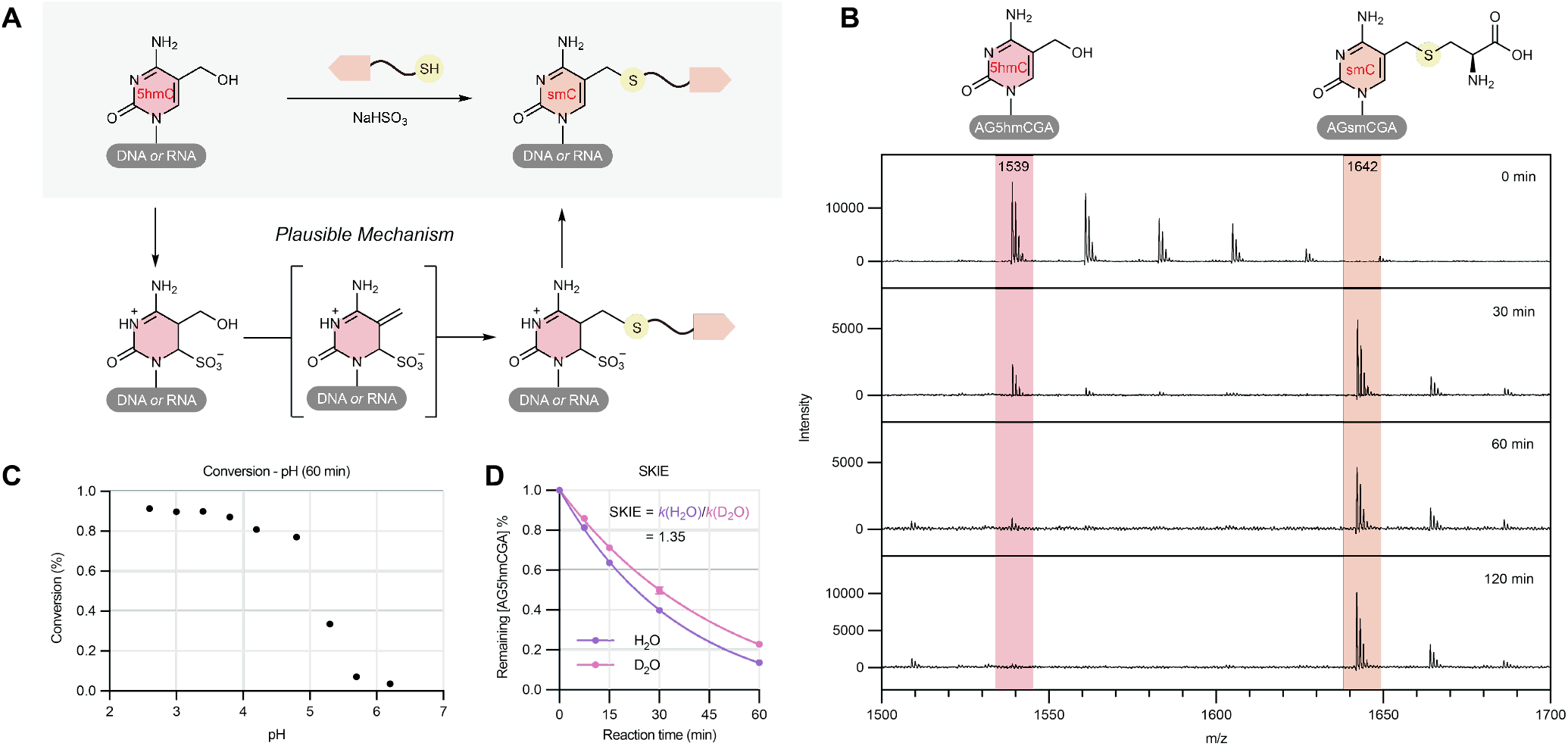
Optimization and mechanistic validation of BALT. (A) Proposed reaction mechanism involving bisulfite-mediated activation of 5hmC to generate a reactive intermediate, followed by nucleophilic thiol addition to yield a stable sulfur-containing adduct (smC). (B) BALT conversion of a DNA probe (5”-AG5hmCGA) with 1 M bisulfite at pH = 3 monitored by MALDI-TOF MS, confirming formation of smC from 5hmC over time. (C) pH-dependent conversion efficiency of 5hmC, showing optimal labeling under mildly acidic conditions. (D) Time-dependent consumption of 5hmC substrate, indicating first-order kinetics and solvent kinetic isotope effect analysis (SKIE = 1.35) supporting proton transfer involvement in the rate-determining step.

To start, we selected a 5hmC-containing DNA oligomer probe (5”-AG5hmCGA) as our model substrate and cysteine as model thiol nucleophile to optimize labeling condition. MALDI-TOF MS analysis showed that upon coincubation with bisulfite and cysteine, the peak (*m*/*z* 1539), corresponding to the starting 5hmC oligonucleotide, decreased, and a new peak (*m*/*z* 1642), corresponding to the thiol-labeled product, appeared (**Figure 2B**), which realized almost full conversion in 2 hours. Under the same condition, unmodified cytosine showed no labeling (**Figure S1**).

During optimization, we observed that the reaction was highly dependent on pH. We investigated the conversion rate of DNA oligomer in 1 hour and found that the reactivity varies quite a bit based on pH values (**Figure 2C**). It is noteworthy that the N3 position of cytosine nucleosides exhibits a p*K*a around 4.1 under aqueous conditions.^31^ Under less acidity, conversion was poor, consistent with inadequate cytosine protonation and reduced formation of the reactive intermediate (**Figure 2A**). A mildly acidic window near pH 3 provided the best balance between conversion, substrate integrity, and suppression of unwanted cytosine deamination. Moreover, the reaction showed time-dependent consumption of 5hmC, and kinetic fitting was consistent with apparent first-order behavior under conditions in which bisulfite and thiol were in large excess (**Figure 2D**). A solvent kinetic isotope effect of approximately 1.35 indicated that proton transfer contributes to the rate-determining process. This mechanistic interpretation is further supported by control experiments. First, replacing bisulfite with acetate at the same pH eliminated productive labeling, showing that acidity alone is not sufficient. Second, omitting thiol shifted the reaction toward CMS, demonstrating that bisulfite is both an activator and a competing nucleophile (**Figure S2**).

A key practical advantage of this chemistry is that it distinguishes 5hmC from 5hmU. 5hmU-containing mismatches can arise through deamination pathways and may be recognized by enzyme-based labeling.^15^ In assays of our synthetic-probe (5”-AG5hmUGA), BALT did not productively label 5hmU under conditions that efficiently labeled 5hmC (**Figure S3**). This selectivity is consistent with the requirement for cytosine-specific activation in the proposed mechanism (**Figure 2A**).

To transform BALT into a facile sequencing method, we synthesized several biotin-thiol reagents through straightforward condensation reaction (**Figure 3A**). The simplest designs linked biotin to cysteamine or cysteine, while additional probes incorporated polyethylene glycol spacer. After screening, we observed clean and nearly complete conversion of 5hmC using biotin-cysteamine in 30 minutes (**Figure 3B**). Its higher reaction rate relative to cysteine likely arises from differences in thiol nucleophilicity under the acidic reaction conditions. The protonated *α*-amino group of cysteine decreases the nucleophilicity of its thiol, whereas the corresponding amino group in biotin– cysteamine is acylated and therefore does not carry a positive charge. We also found that Biotin-PEG-SH improved solubility but produced unexpected fragment ion signals. In addition, a previously reported similar probe, biotin-cysteine,^29^ differs from biotin-cysteamine by only one extra carboxylic acid group, but it showed very poor water solubility and generated almost no expected 5hmC adduct under our reaction conditions. Using 5hmC and 5hmU nucleosides as model substrates, we observed quantitative conversion of 5hmC to the biotin-thioether product (bsmC) under the biotin-labeling conditions (**Figure S4**), whereas no detectable reaction was observed for 5hmU (**Figure S5**). These results further demonstrate that the chemical labeling strategy can selectively distinguish 5hmC from 5hmU.

**Figure 3.**
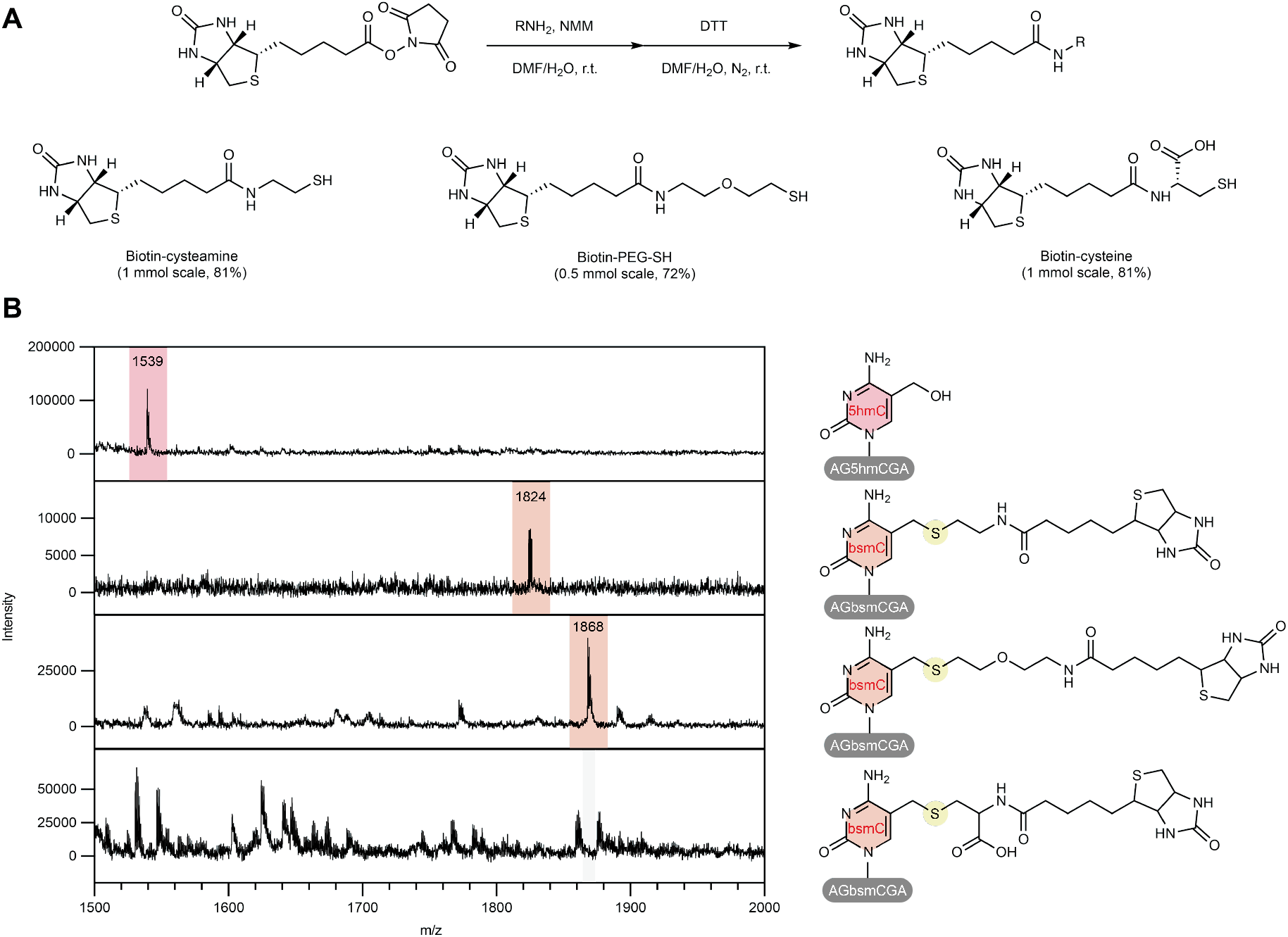
Synthesis and screening of biotin-thiol reagent to realize one-step labeling. (A) Facile synthesis route of biotin-thiol probes. (B) Biotin-thiol probes incorporated into the BALT reaction, in which biotin-cysteamine shows clean conversion and efficient formation of the desired labeled product.

We next used a 164-nt oligonucleotide containing a single 5hmC site to evaluate whether DNA polymerases could efficiently bypass the resulting bsmC adduct. Primer-extension assays showed that full-length extension products predominated across the DNA polymerases tested, indicating efficient read-through of bsmC with minimal polymerase stalling. This behavior contrasts with that reported for 5hmC-Seal, in which the bulky side-chain adduct causes notable *Taq* polymerase stalling and a large portion of truncated extension products.^13^ To evaluate the polymerase readout following BALT chemistry, we performed Sanger sequencing and examined whether bsmC could resist APOBEC-mediated deamination, thereby enabling single-nucleotide-resolution detection. Under mild bisulfite conditions, no appreciable C-to-U deamination was observed, and 5hmC remained read as C (**Figure S6**). In contrast, when BALT labeling was coupled with APOBEC deamination, labeled 5hmC was retained as C, whereas unmodified C and 5mC were converted to U. These results indicate that BALT chemistry is not limited to enrichment-based profiling but can also support higher-resolution mapping through the combination of biotin enrichment, three-letter sequencing, and an appropriate mapping strategy.

The sequencing workflow was then designed with a single chemical labeling step (**Figure 4A**). Genomic DNA was fragmented, end-repaired, and ligated to sequencing adapters before BALT treatment. Adapter-ligated fragments were exposed to bisulfite and biotin-cysteamine, enriched with streptavidin beads, amplified by PCR, and sequenced. This order preserves library molecules after enrichment and avoids multiple post-labeling manipulations. Gel and PCR amplification assays showed that the optimized treatment caused limited degradation (**Figure S8**) and allowed efficient PCR amplification (**Figure S9**). Using equal amounts of input DNA, the BALT workflow required fewer PCR amplification cycles than 5hmC-Seal when *Taq* polymerase was used. This improvement is consistent with previous reports that the bulky adduct generated by 5hmC-Seal can cause steric hindrance and polymerase stalling.^13^ In addition, BALT incorporates a desulfonation step after biotin labeling, which converts residual sulfonated cytosine byproducts back to cytosine and thereby improves polymerase read-through and overall amplification efficiency. We incorporated an equimolar mixture of 5hmC-and non-5hmC-containing spike-in controls at a 1:1 input ratio. Following labeling and enrichment, the 5hmC-containing spike-in reads exhibited a 28.6-fold enrichment relative to the non-5hmC-containing controls (**Figure S10**), demonstrating the high efficiency and selectivity of the BATL workflow at the sequencing-library level.

**Figure 4.**
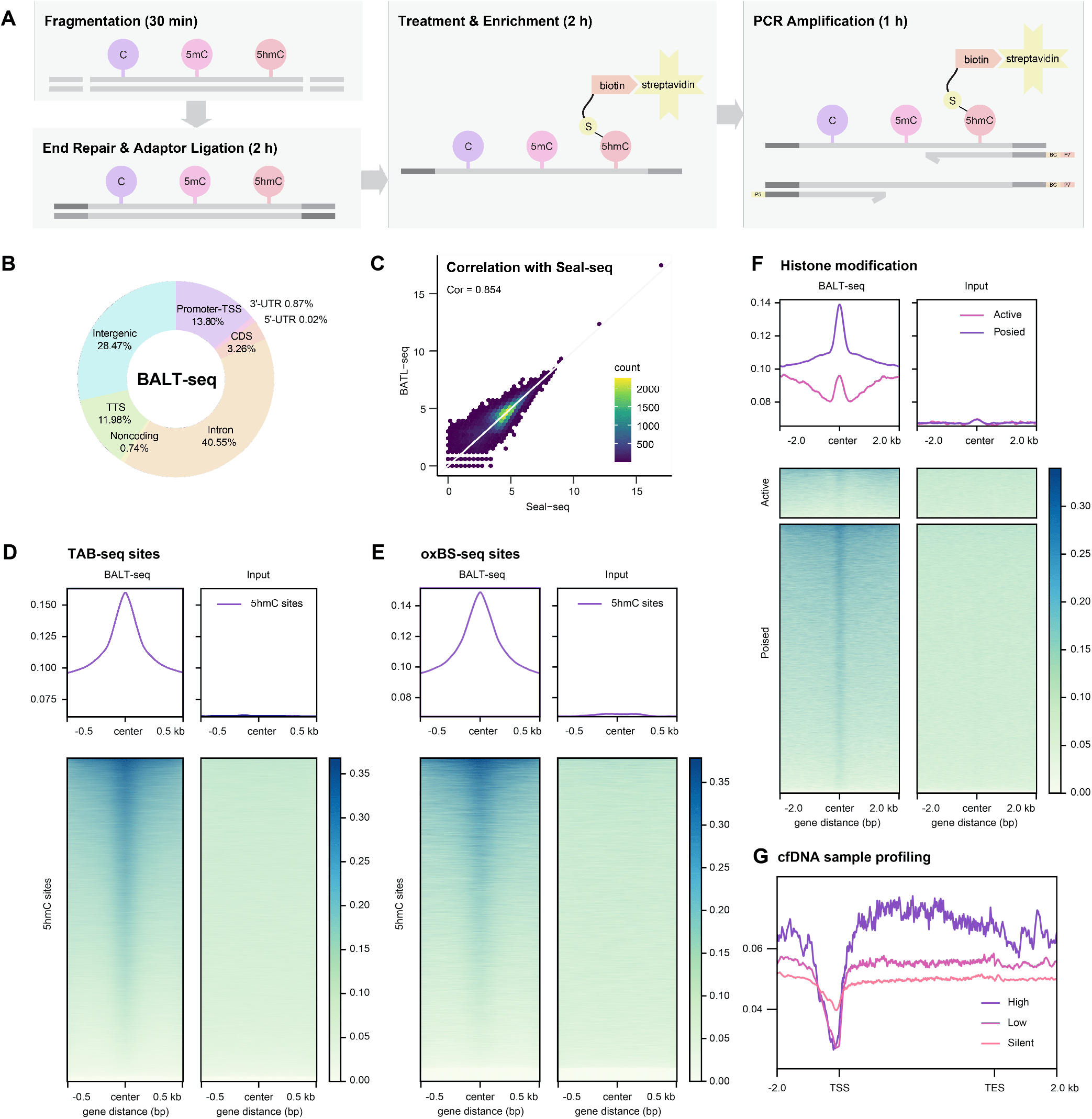
Workflow and genome-wide performance of DNA BALT-seq. (A) Experimental workflow of BALT-seq, including DNA fragmentation, adaptor ligation, bisulfite-thiol treatment, enrichment via biotin-streptavidin interaction, and PCR amplification. (B) Genomic annotation of BALT-seq peaks, showing distribution across intergenic, intronic, promoter, and coding regions. (C) Correlation of BALT-seq with Seal-seq, demonstrating high concordance (Pearson correlation = 0.854). (D) Metagene analysis of BALT-seq signal over 5hmC sites identified by TAB-seq. (E) Metagene analysis of BALT-seq signal over 5hmC sites identified by oxBS-seq. (F) Distribution of BALT-seq signals relative to histone modification states (active versus poised chromatin), showing preferential enrichment in poised regions. (G) Profiling of cfDNA samples, revealing distinct 5hmC patterns across transcriptional states (high, low, silent).

BALT-seq was then applied to mouse embryonic stem cell (mESC) genomic DNA (gDNA) to evaluate genome-wide performance. Peak annotation uncovered BALT-seq signals predominantly across intronic, intergenic, and promoter regions (**Figure 4B**). The pattern is expected for 5hmC, which marks both gene bodies and distal regulatory elements. Side-by-side comparison with 5hmC-Seal produced strong concordance, with a Pearson correlation coefficient of 0.854 and extensive overlap between called peaks (**Figure 4C**). We next compared mESCs gDNA BALT-seq profiles with base-resolution 5hmC maps generated by TET-assisted bisulfite sequencing (TAB-seq)^32^ (**Figure 4D**) and oxidative bisulfite sequencing (oxBS-seq)^33^ (**Figure 4E**). TAB-seq uses enzymatic protection and oxidation to reveal 5hmC at single-base resolution, while oxBS-seq chemically oxidizes 5hmC before bisulfite conversion to derive 5hmC by comparison with conventional bisulfite sequencing. BALT-seq signal was enriched over TAB-seq-defined and oxBS-seq-defined 5hmC sites, while input controls showed little enrichment. Agreement with enzymatic glycosylation-click strategy and two base-resolution methods indicates that BALT-seq peaks correspond to genuine 5hmC-enriched loci.

The biological patterns recovered by BALT-seq are also consistent with established 5hmC functions. Previous genomic studies have shown that 5hmC is enriched in gene bodies and enhancers and is associated with transcriptional regulation in mammalian cells and human tissues.^4,5,10^ In our data, BALT-seq signal is elevated over gene bodies and correlated with transcriptionally active chromatin states. Analysis relative to histone modifications further showed that BALT-seq peaks preferentially mark poised chromatin features, including H3K4me1-positive regions with limited H3K27ac signal, rather than strongly active enhancers bearing both H3K4me1 and H3K27ac (**Figure 4F**). This pattern is consistent with reports that oxidized cytosine derivatives can mark regulatory states linked to enhancer priming and active demethylation intermediates.^34^

The mildness of BALT chemistry suggested potential application to fragmented and low-input samples. Circulating cell-free DNA is an especially demanding substrate because it is short, dilute, and easily lost during multi-step processing. Plasma cfDNA typically shows a nucleosomal fragment distribution centered near 167 bp, while urinary cfDNA can be substantially shorter.^31^ Harsh bisulfite conversion can destroy much of this material, making classical base-conversion methods challenging to many liquid biopsy samples. In contrast, BALT operates under low-degradation conditions. Building on earlier studies that classified genes into three groups based on cfRNA expression,^32^ we plotted the average cell-free 5hmC distribution across gene bodies and observed pronounced enrichment in and around the gene bodies of highly expressed genes (**Figure 4G**). Consistent with previous reports, these results demonstrate that cell-free 5hmC originates from multiple tissue types and retains information beyond that derived from blood, which may serve as potential disease biomarkers.^7,10,37^ BALT-seq shortens the library workflow, reduces purification steps, and avoids expensive enzymatic labeling reagents. Its simplicity and robustness offer distinct advantages across plasma, urine, and disease contexts.

Because BALT strategy is completely chemistry based without relying on any enzymatic activity, we next extended this approach to RNA. RNA hm^5^C and hm^5^C*m* contain the same hydroxymethylcytosine base as DNA 5hmC, but RNA is prone to hydrolysis, and is analyzed through reverse transcription followed by polymerase amplification. In addition, hm^5^C/hm^5^C*m* abundance in RNA is low compared with canonical ribonucleosides. Earlier LC-MS work demonstrated formation of hm^5^C from m^5^C in RNA and identified hm^5^C*m* as a second oxidative derivative, but transcriptome-wide chemical enrichment methods remain underdeveloped.^18,19^ The recent demonstration that RNA m^5^C oxidation can regulate chromatin state through retrotransposon RNA further begs for methods that can map oxidized RNA cytosines.^20^

A synthetic hm^5^C-containing RNA oligomer probe (5”-AGhm^5^CGA) was therefore subjected to BALT conditions and monitored by MALDI-TOF MS (**Figure 5A**). Compared with DNA 5hmC, RNA hm^5^C was less reactive and approached an apparent plateau after treatment for one hour. The reduced conversion (66 %) may reflect altered hydrogen bond system, or differences in sugar conformation that modulate reactivity of the cytosine ring. Nevertheless, the conversion was sufficient for enrichment.

**Figure 5.**
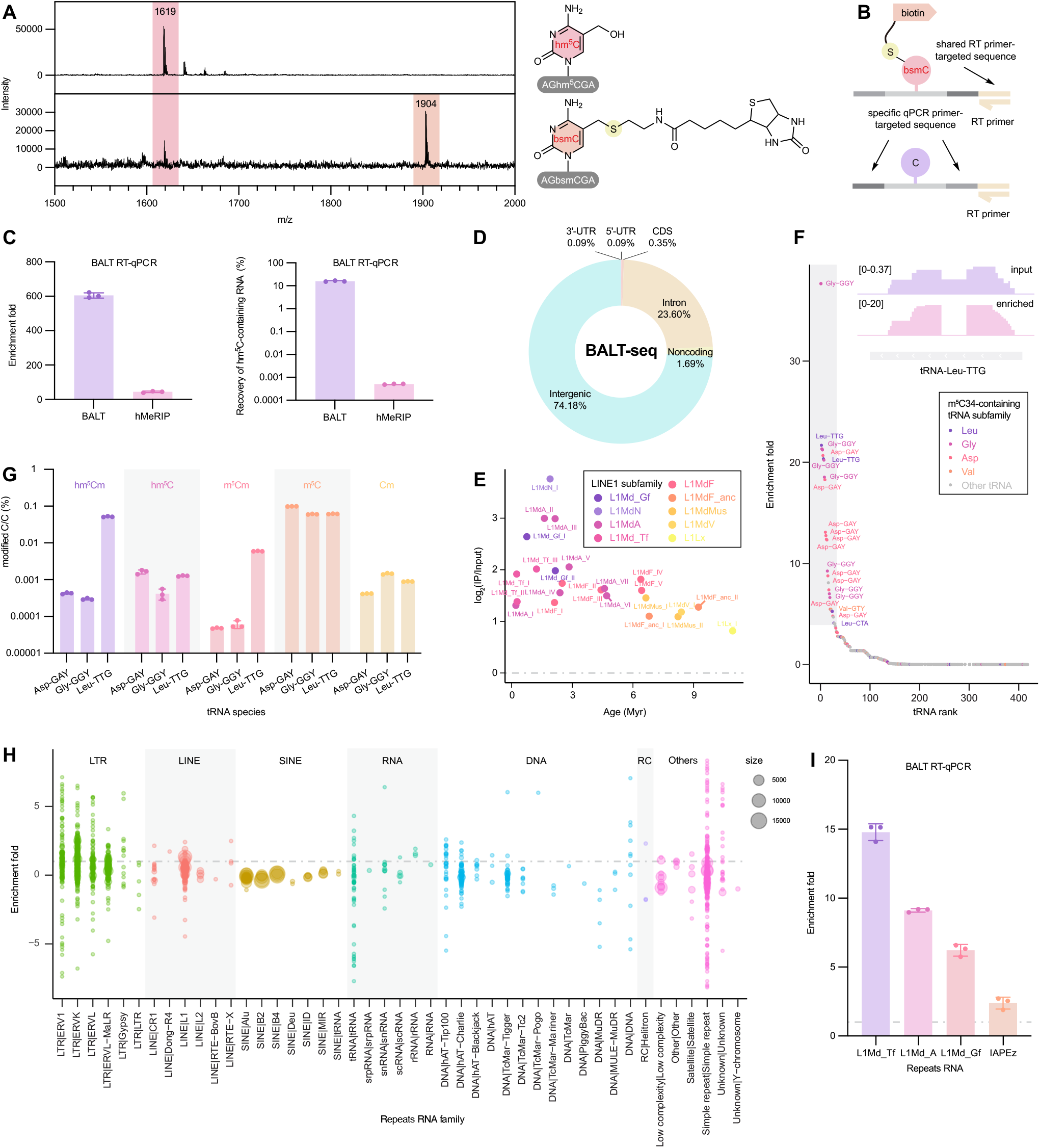
Application of BALT to RNA hm^5^C/hm^5^C*m* profiling and transcriptomic features. (A) BALT conversion of an RNA probe (5”-AGhm^5^CGA) monitored by MALDI-TOF MS after treatment for 1 h. (B) Schematic of RT-qPCR strategy for validation of enrichment of labeled hm^5^C-containing RNA. (C) Comparison of enrichment and recovery efficiency between BALT and hMeRIP, showing higher enrichment and recovery rate by BALT. (D) Genomic annotation of RNA BALT-seq peaks, showing predominant mapping to intergenic and intronic regions. (E) Relationship between evolutionary age and hm^5^C/hm^5^C*m* enrichment across mouse LINE1 subfamilies. (F) Enrichment ranking of tRNA species, indicating selective modification patterns among tRNA isoacceptors. (G) Validation and quantification of modified cytosine fractions (hm^5^C*m*, hm^5^C, m^5^Cm, m^5^C and C*m*) across different identified hm^5^C/hm^5^C*m*-containing tRNA species. (H) Global enrichment landscape across repeat RNA families, illustrating differential hm^5^C/hm^5^C*m* patterns among LTR, LINE, SINE, and other elements. (I) RT-qPCR validation of BALT enrichment for representative repeat RNAs, including the LINE1 subfamilies L1Md_Tf, L1Md_A, and L1Md_Gf and the endogenous retroviral element IAPEz.

Next, we designed two *in vitro*-transcribed RNA oligonucleotides: one containing 10% hm^5^C and another matched control lacking hm^5^C. These two RNAs share the same reverse transcription (RT) primer-targeted sequence but contain different qPCR primer-targeted regions. Using 100 ng of RNA input, we validated enrichment with an RT-qPCR strategy in which labeled hm^5^C-containing RNA was captured by streptavidin beads and quantified relative to non-hm^5^C-containing input controls (**Figure 5B**). BALT enrichment outperformed hMeRIP in both enrichment magnitude and recovery (**Figure 5C**). This result is not surprising that hMeRIP relies on antibody recognition of a small modified nucleoside embedded within structured DNA or RNA, whereas BALT converts the modification into a covalent biotin handle, providing a much stronger and more selective binding than antibody affinity or epitope accessibility.

The global abundance of hm^5^C/hm^5^C*m* in mouse embryonic stem cell RNA was compatible with an enrichment-based workflow which motivated us to perform RNA BALT-seq on total RNA. Genomic annotation of RNA BALT-seq peaks showed a dominant fraction assigned to intergenic regions, with substantial intronic signal and smaller contributions from annotated noncoding and coding features (**Figure 5D**), which suggests that hm^5^C/hm^5^C*m* may be enriched in noncanonical or repeat-associated RNA populations rather than primarily in conventional mRNA exons.

Repeat-family analysis supported this interpretation. BALT-seq signal was not evenly distributed across repetitive elements; instead, specific families showed stronger enrichment, including LINE and LTR-associated classes (**Figure 5H**) such as L1Md_T, L1Md_A, and IAPEz-int (**Figure S11**), which is further validated by BALT RT-qPCR (**Figure 5I**). In LINE1 subfamily, we observed that hm^5^C/hm^5^C*m* is preferentially enriched on evolutionarily young LINE1 RNAs (**Figure 5E**), suggesting that hm^5^C/hm^5^C*m* may mark transcriptionally competent repeat RNAs for nuclear regulation. Because repetitive RNAs can regulate chromatin and developmental gene expression, BALT-seq may help future studies on how methylcytosine oxidation in RNA may regulate chromatin state during various physiological processes.

A second prominent feature of RNA BALT-seq was enrichment of specific tRNA species. tRNAs are among the most heavily modified RNA classes, and cytosine modifications at or near the anticodon loop and variable regions can influence decoding, stability, and stress responses.^38,39^ Interestingly, although multiple m^5^C sites have been reported on tRNAs, only the m^5^C34-containing tRNAs (Leu, Gly, Asp and partial Val tRNAs)^40^ showed strong enrichment in RNA BALT-seq (**Figure 5F**). Because hm^5^C and hm^5^C*m* are oxidative derivatives of methylated cytosines, selective tRNA enrichment may reflect regulated oxidation of pre-existing m^5^C-or m^5^C*m*-modified cytosines. Notably, conventional bisulfite sequencing cannot distinguish m^5^C/m^5^C*m* from hm^5^C/hm^5^C*m*, raising the possibility that a subset of sites previously assigned as m^5^C/m^5^C*m* may actually represent hm^5^C/hm^5^C*m*, even if they constitute a minor fraction. Among the enriched tRNAs, tRNA-Leu-TTG was previously reported to harbor hm^5^C*m* in mature tRNA.^37^ We observed approximately 50-fold enrichment together with a pronounced spliced-read pattern, consistent with a biogenesis model in which tRNA splicing precedes the proposed ALKBH1-mediated oxidation.^17,41^

Targeted validation of modified cytosine fractions across abundant candidate tRNAs further supported differential modification (**Figure 5G**). The relative abundances of hm^5^C*m*, hm^5^C, m^5^C*m*, m^5^C, and C*m* varied across the enriched tRNA species.

The coexistence of methylated, hydroxymethylated, and ribose-methylated derivatives suggests that hm^5^C/hm^5^C*m* formation is carefully regulated. Although tRNA-Leu-TTG showed the highest hm^5^C*m* abundance, other m m^5^C34-containing tRNAs also appeared to undergo oxidation and ribose methylation to varying extents, potentially through a shared enzymatic pathway.

## CONCLUSION

Over the past decade and a half, the development of 5hmC mapping technologies has substantially advanced understanding of hydroxymethylcytosine distribution and function across mammalian genomes. Early antibody-based and anti-CMS enrichment approaches enabled genome-wide detection of 5hmC, while *β*-glucosyltransferase-based selective labeling, exemplified by 5hmC-Seal, provided sensitive and robust profiling applicable to increasingly limited biological material.^12–14,27,28^ In parallel, TAB-seq and oxBS-seq established single-nucleotide-resolution measurement of 5hmC and helped define its characteristic enrichment in gene bodies and regulatory regions.^4,5,32,33^ These technologies have collectively revealed tissue-specific 5hmC landscapes and demonstrated the diagnostic potential of 5hmC signatures in circulating cell-free DNA.^7^ However, existing approaches achieve sensitivity and resolution through distinct combinations of antibody recognition, enzymatic derivatization, or chemical/base-conversion workflows, leaving room for simpler chemoselective strategies that directly functionalize hydroxymethylcytosine in both DNA and RNA.

BALT-seq establishes a chemoselective route to hydroxymethylcytosine enrichment that is conceptually distinct from antibody recognition, enzymatic glycosylation, and base-conversion sequencing. By tuning bisulfite chemistry to activate the 5-hydroxymethyl group while suppressing canonical cytosine deamination, BALT converts 5hmC, hm^5^C, and hm^5^C*m* into covalently biotinylated products suitable for streptavidin enrichment. Mechanistic experiments support a competitive nucleophilic capture model in which thiol addition competes with CMS formation.

In genomic DNA, BALT-seq produces 5hmC maps that agree well with 5hmC-Seal-seq, TAB-seq, and oxBS-seq while reducing reliance on enzymes and tedious procedures. It recovers expected gene-body and chromatin-state distributions in mouse embryonic stem cells and is compatible with low-degradation cfDNA analysis. These features make BALT-seq suitable for epigenomic discovery, low-input profiling, and potentially clinical-scale 5hmC biomarker studies.

The extension to RNA expands the scope of hydroxymethylcytosine biology. RNA BALT-seq enriches hm^5^C/hm^5^C*m*-containing molecules more effectively than antibody-based enrichment and reveals structured modification patterns across tRNA species and other repeat-associated RNA families. These results provide a foundation for studying oxidized RNA cytosines in development, chromatin regulation, and disease. More broadly, BALT-seq shows that classical bisulfite chemistry, long used only to read cytosine modifications, can be redirected to functionalize cytosine modifications in nucleic acids.

## Supporting information

Supplementary figures, materials and methods and NMR spectra

## ASSOCIATED CONTENT

### Data Availability Statement

All sequencing data are available at the GEO database (accession: GSE341613, GSE341614, GSE 341615). All relevant additional data have been published with the manuscript, either as part of the main text or in the supplement.

### Supporting Information

The Supporting Information is available free of charge on the ACS Publications website.

Supplementary figures, materials and methods and NMR spectra (PDF)

## AUTHOR INFORMATION

### Authors

Jiahao Li – *Department of Chemistry, Department of Biochemistry and Molecular Biology, Institute for Biophysical Dynamics, The University of Chicago, Chicago, IL, USA; Howard Hughes Medical Institute, The University of Chicago, Chicago, IL, USA*.

Pei-Hong Zhang – *Department of Chemistry, Department of Biochemistry and Molecular Biology, Institute for Biophysical Dynamics, The University of Chicago, Chicago, IL, USA; Howard Hughes Medical Institute, The University of Chicago, Chicago, IL, USA*.

Yanchu Arvin Wang – *Department of Chemistry, Department of Biochemistry and Molecular Biology, Institute for Biophysical Dynamics, The University of Chicago, Chicago, IL, USA; Howard Hughes Medical Institute, The University of Chicago, Chicago, IL, USA*.

Yuhao Zhong – *Department of Chemistry, Department of Biochemistry and Molecular Biology, Institute for Biophysical Dynamics, The University of Chicago, Chicago, IL, USA; Howard Hughes Medical Institute, The University of Chicago, Chicago, IL, USA*.

Yiding Wang – *Department of Chemistry, Department of Biochemistry and Molecular Biology, Institute for Biophysical Dynamics, The University of Chicago, Chicago, IL, USA; Howard Hughes Medical Institute, The University of Chicago, Chicago, IL, USA*.

Ruitu Lyu – *Department of Chemistry, Department of Biochemistry and Molecular Biology, Institute for Biophysical Dynamics, The University of Chicago, Chicago, IL, USA; Howard Hughes Medical Institute, The University of Chicago, Chicago, IL, USA*.

Chenyou Zhu – *Department of Chemistry, Department of Biochemistry and Molecular Biology, Institute for Biophysical Dynamics, The University of Chicago, Chicago, IL, USA; Howard Hughes Medical Institute, The University of Chicago, Chicago, IL, USA*.

### Author Contributions

C. H., Q. D., and J. L. conceived the study. J. L., with assistance from Y. A. W. and contribution from Y. Z., Y. W., R. L., C. Z., performed reaction optimization and library preparation. P.-H. Z. analyzed the sequencing data. J. L. and C. H. wrote the paper with inputs from all authors. All authors discussed the results and commented on the paper.

### Notes

The authors declare following competing financial interest(s). C.H. is a scientific founder, a member of the scientific advisory board and equity holder of AllyRNA, Inc., and a scientific cofounder and equity holder of Accent Therapeutics, Inc., and Ellis Bio, Inc.. The other authors declare no competing interests. C. H., Q, D,, and J. L. are on a patent filed regarding hydroxymethylcytosine labeling and sequencing using BALT-seq.

## ACKNOWLEDGMENT

We are grateful to the University of Chicago Genomic Facility for assistance with high-throughput sequencing. This work is supported by the National Institute of Health R01 HG006827. C. H. is an investigator of the Howard Hughes Medical Institute

