## Supplementary figures, materials and methods and NMR spectra for "A One-Step Chemoselective Strategy for Hydroxymethylcytosine Sequencing in DNA and RNA"

† These authors contributed equally.

### Supplementary Figures

S1

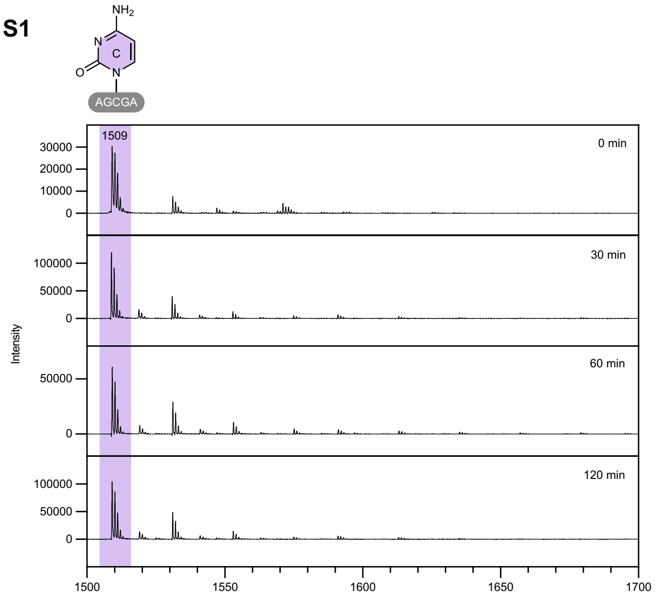


S2

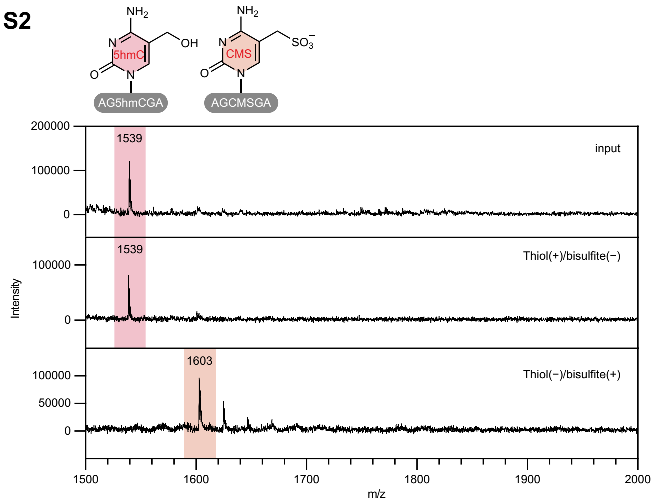


S3

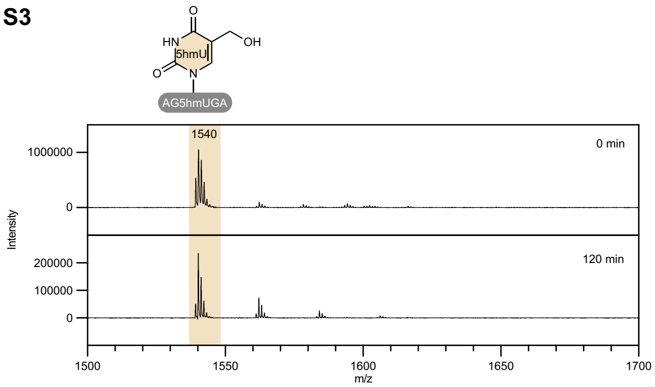


S4

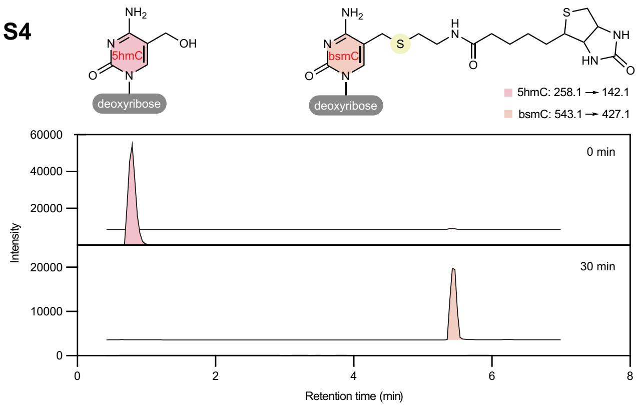


S5

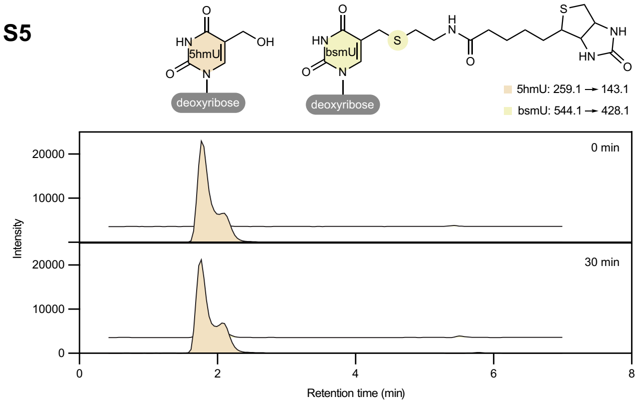

S6

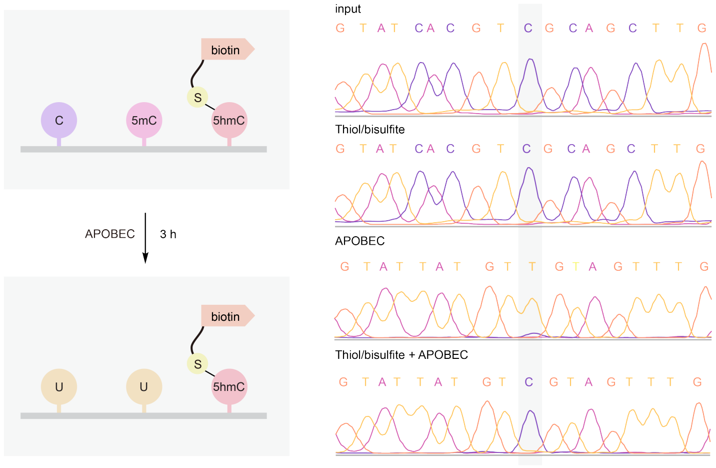


S7

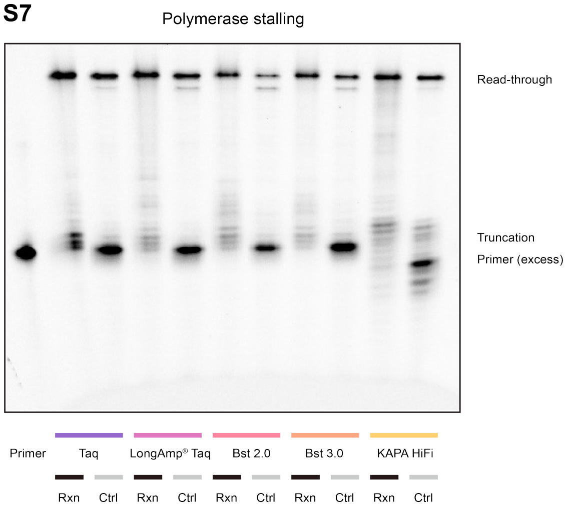


S8

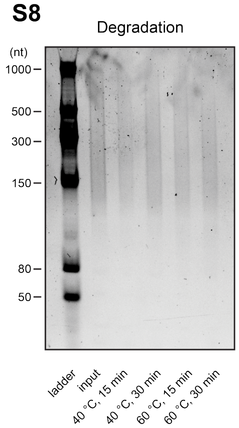


S9

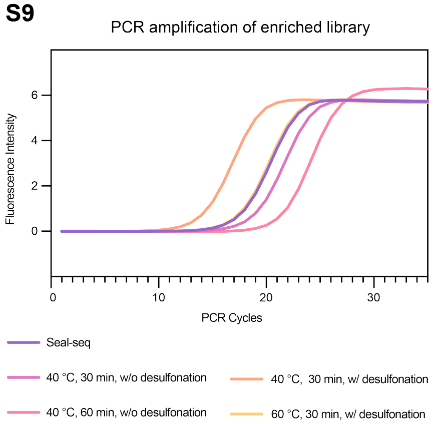


S10

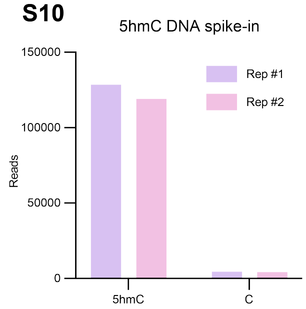


S11

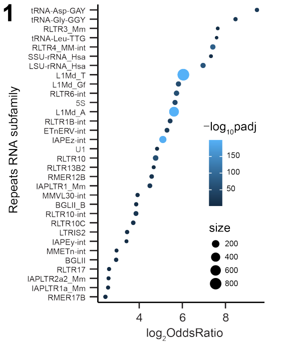


**Figure S1. Selectivity of BALT toward 5hmC over unmodified cytosine.**

MALDI-TOF MS analysis of an unmodified cytosine-containing DNA probe (5′-AGCGA) under BALT reaction conditions for 0, 30, 60, and 120 min. No detectable thiol-labeled product was observed, demonstrating minimal reactivity of unmodified cytosine under the optimized conditions.

**Figure S2. Control experiments supporting bisulfite-mediated activation of 5hmC.**

MALDI-TOF MS analysis of a 5hmC-containing DNA probe (5′-AG5hmCGA) under control conditions. In the absence of bisulfite, addition of thiol did not induce detectable conversion of 5hmC. In the absence of thiol, bisulfite treatment converted 5hmC predominantly to the cytosine-5-methylenesulfonate (CMS) product, supporting a competitive nucleophilic capture mechanism between thiol and bisulfite.

**Figure S3. Selectivity of BALT toward 5hmC over 5hmU.**

MALDI-TOF MS analysis of a 5hmU-containing DNA probe (5′-AG5hmUGA) before and after 120 min of BALT treatment. No detectable thiol-labeled product was observed, demonstrating that 5hmU is not reactive under conditions that efficiently label 5hmC.

**Figure S4. Efficient biotin labeling of 5hmC nucleoside by BALT.**

LC-MS/MS analysis of 5hmC nucleoside before and after 30 min of BALT treatment with biotin-cysteamine. Disappearance of the 5hmC signal and formation of the corresponding biotin-thioether product (bsmC) demonstrate rapid and near-quantitative labeling of 5hmC.

**Figure S5. Lack of detectable biotin labeling of 5hmU nucleoside by BALT.**

LC-MS/MS analysis of 5hmU nucleoside before and after 30 min of BALT treatment with biotin-cysteamine. No detectable formation of the corresponding biotin-thioether product (bsmU) was observed, further demonstrating the chemical selectivity of BALT for hydroxymethylcytosine over hydroxymethyluracil.

**Figure S6. Compatibility of BALT-labeled 5hmC with APOBEC-mediated three-letter sequencing.**

Schematic and Sanger sequencing of DNA containing C, 5mC, and 5hmC following BALT labeling and/or APOBEC treatment. BALT treatment alone preserved cytosine identity, whereas APOBEC treatment converted unmodified C and 5mC to U, read as T during sequencing. In contrast, BALT-labeled 5hmC remained protected from APOBEC-mediated deamination and was retained as C, supporting the potential application of BALT to base-resolution 5hmC detection.

**Figure S7. Polymerase read-through of BALT-labeled 5hmC.**

Primer-extension analysis of a 164-nt DNA template containing a single 5hmC site following BALT reaction (Rxn) or control treatment (Ctrl). Extension products generated by *Taq*, LongAmp® *Taq*, *Bst* 2.0®, *Bst* 3.0®, and KAPA HiFi DNA polymerases are shown. Full-length read-through products predominated across the tested polymerases, indicating limited polymerase stalling induced by the BALT-generated bsmC adduct.

**Figure S8. Evaluation of DNA integrity under BALT reaction conditions.**

Gel electrophoresis analysis of DNA before and after BALT treatment at 40 °C or 60 °C for 15 or 30 min. The optimized mild reaction conditions preserved the majority of input DNA and caused limited degradation.

**Figure S9. PCR amplification efficiency of BALT-enriched DNA libraries.**

PCR amplification curves of enriched DNA libraries prepared using 5hmC-Seal or BALT under the indicated reaction conditions. Incorporation of a desulfonation step improved amplification efficiency, with BALT-treated libraries requiring fewer amplification cycles than the 5hmC-Seal library under the optimized condition.

**Figure S10. Selective enrichment of 5hmC-containing DNA spike-in controls by BALT-seq.**

Sequencing read counts of 5hmC-containing and control DNA spike-ins following BALT labeling and streptavidin enrichment in two independent replicates. Strong preferential recovery of the 5hmC-containing spike-in demonstrates efficient and selective enrichment at the sequencing-library level.

**Figure S11. Enrichment of repeat RNA subfamilies identified by RNA BALT-seq.**

Enrichment analysis of repeat RNA subfamilies in mESC RNA BALT-seq. Strongly enriched repeat subfamilies include LINE- and LTR-associated elements, consistent with preferential hm5C/hm5Cm enrichment in specific repeat-derived RNA populations.

#### Materials and Methods

##### General BALT treatment of model DNA and RNA

Prepare 2 M sodium bisulfite solution (2× SBS) by dissolving 190 mg sodium metabisulfite ($Na_2S_2O_5$; Sigma-Aldrich, cat. no. S9000) in nuclease-free water and adjusting the final volume to 1 mL. Vortex vigorously until completely dissolved, add 37% hydrochloric acid (HCl; Sigma-Aldrich, cat. no. 320331) to adjust pH as required and vortex again for 5 seconds. Denature substrate DNA by incubating in a preheated thermal cycler for 5 minutes at 95 °C (for RNA, 70 °C) with heated lid set to 105 °C. Transfer tube to ice immediately.

Condition 1: Cysteine labeling

Prepare 2.5 M cysteine solution (2× Cys) by dissolving 302 mg cysteine (Sigma-Aldrich, cat. no. 168149) in nuclease-free water and adjusting the final volume to 1 mL Vortex vigorously until completely dissolved. Add 2× SBS and 2× Cys at a 1:1 volume ratio. Assemble the mixture with the model DNA or RNA and incubate in a preheated thermal cycler as required. (Optimized condition for short oligonucleotides: 60 minutes at 40 °C)

Condition 2: Biotin-thiol labeling

Weigh 2 mg biotin-thiol in a 200 µL tube and add 25 µL 2× SBS solution and preheat at 80 °C for 5 min for saturation of thiol. Assemble the saturated supernatant with the model DNA or RNA and in a preheated thermal cycler as required. (Optimized condition for short oligonucleotides: 60 minutes at 40 °C)

##### Preparation of model DNA and RNA for MALDI-TOF MS

Regular and 5hmC/hm$^5$C-labeled short oligonucleotides for MALDI-TOF MS assay were purchased from IDT. 5hmU-labeled short oligonucleotide 5'-AG5hmUGA for MALDI-TOF MS assay was prepared by APOBEC3A treatment (New England Biolabs) for 3 h according to manufacturer's manual starting from 5'-AG5hmCGA.

##### MALDI-TOF MS assay

The Matrix Assisted Laser Desorption Ionization Time of Flight Mass Spectrometry (MALDI-TOF MS) assay was performed treating synthetic oligonucleotides under general BALT treatment protocol in 20 µL scale. Afterwards, to remove sodium ion, the reaction mixture was treated by ion exchange resin AmberChrom™, 50WX8 200-400 (H) (Thermo Fisher Scientific). Then, 1.6 µL of the mixture was combined with 1.6 µL of a matrix consisting of 2,4,6-trihydroxyacetophenone (THAP) monohydrate on a MALDI plate. The MALDI-TOF MS spectrum was recorded on a Bruker Ultra-flex TOF/TOF MALDI mass spectrometer using the negative ion reflection mode. Data was processed in the Bruker Flex Analysis software 3.4.

##### Preparation of model DNA for primer extension

5hmC-labeled 82-mer oligonucleotides for primer extension assay were purchased from IDT.

Template:
GTGACTGGAGTTCAGACGTGTGCTCTGCCTCCGATCTAGATGTGTAGTATCACGT5hmCGCAGCTTGACC
GCTCTAGTGACGGCT

Primer: 5'-FAM-AGCCGTCACTAGAGCGGTCAAGC

##### Primer extension assay

100 ng of 82-mer DNA template was treated under general BALT treatment protocol in 20 µL scale using biotin-cysteamine. After the treatment, ssDNA was desalted using a Bio-Spin P-6 Gel Column (Bio-Rad) and purified by Oligo Clean & Concentrator (Zymo Research), and 40 ng purified ssDNA was used for primer extension reaction. 1 µL of 1 µM FAM-primer was added to treated ssDNA and annealed at 65 °C for 5 min, then put on ice for 2 min. Then,

polymerase 2× buffer were added and incubated according to manufacturer's protocol including *Taq* (New England Biolabs), LongAmp® *Taq* (New England Biolabs), *Bst* 2.0® (New England Biolabs), *Bst* 3.0® (New England Biolabs) and KAPA HiFi (Roche). After reaction, 5 µL of the reaction mixture was incubated with 5 µL 2× RNA loading dye (New England Biolabs) and incubated at 95 °C for 5 min and immediately load onto 15% denaturing polyacrylamide gel and run at 180 V for 90 min. Then, the gel was imaged by Bio-Rad ChemiDoc Imager.

**Sanger sequencing**

100 ng of 82-mer DNA template was treated under general BALT treatment protocol in 20 µL scale. After the treatment, ssDNA was desalted using a Bio-Spin P-6 Gel Column (Bio-Rad) and purified by Oligo Clean & Concentrator (Zymo Research). 16 µL Biotin labeled or unlabeled DNA was mixed with 4 µL formamide, denatured at 85 °C for 10 min, and immediately transferred to a prechilled metal block on ice for approximately 2 min. The denatured DNA was combined on ice with 68 µL nuclease-free water, 10 µL APOBEC Reaction Buffer, 1 µL BSA, and 1 µL APOBEC, which are provided in NEBNext EM-seq kit (New England Biolabs) in a final reaction volume of 100 µL. After thorough mixing, the reaction was incubated at 37 °C for 3 h, purified by Oligo Clean & Concentrator (Zymo Research). Primers matching the 5′ and 3′ flanking regions of the oligo with sanger sequencing primer targeted sequence were used for a ten round of PCR amplification, and the PCR product was used for Sanger sequencing with reverse primer.

Template:
GTGACTGGAGTTCAGACGTGTGCTCTGCCTCCGATCTAGATGTGTAGTATCACGT5hmCGCAGCTTGACC
GCTCTAGTGACGGCT

Reverse Primer: AGCCGTCACTAGAGCGGTCAAGC

Reverse Primer for APOBEC: AACCATCACTAAAACAATCAAAC

Reverse Primer with Sanger Sequencing Primer:

TGTCACCTCCACCCTCTCACTACCACAGCTTCACTCTTCAGCCGTCACTAGAGCGGTCAAGC

Reverse Primer for APOBEC with Sanger Sequencing Primer:

TGTCACCTCCACCCTCTCACTACCACAGCTTCACTCTTCAACCATCACTAAAACAATCAAAC

Forward Primer: GTGACTGGAGTTCAGACGTGTGCTCTG

Forward Primer for APOBEC: GTGATTGGAGTTTAGATGTGTGTTTTG

Sanger Sequencing Primer: TGTCACCTCCACCCTCTC

**DNA degradation test**

100 ng of fragmented mESCs genomic DNA was treated under general BALT treatment protocol in 20 µL scale. After the treatment, ssDNA was desalted using a Bio-Spin P-6 Gel Column (Bio-Rad) and purified by Oligo Clean & Concentrator (Zymo Research). Samples were eluted in 10 µl of water and mixed with 2 µl of Gel Loading Dye Blue (6×) to run on Novex™ TBE Gels 4–20% (Invitrogen) at constant 180 V for 60 min. The gel was stained with 2 µl of SYBR Gold Nucleic Acid Gel Stain (Invitrogen) in 20 ml of TBE buffer at room temperature for 15 min. Then, the gel was imaged by Bio-Rad ChemiDoc Imager.

**tRNA hybrid capture[1]**

Hybrid capture of tRNA was performed with 100 µg DNase-treated, total RNA from mESCs. The hybridization step was done with 200 pmol biotinylated antisense DNA probes (ASO) in 1× SSC buffer by incubation at 95 °C for five minutes and cooling to 22 °C with −0.1 °C/sec temperature ramp. Dynabeads MyOne Streptavidin C1 (Invitrogen) were washed three times in wash buffer (final concentration: 5 mM Tris HCl pH 7.5, 0.5 mM EDTA, 1 M NaCl) once in 5× SSC buffer, and incubated with RNA:DNA hybrid samples in the presence of RNase inhibitor at room temperature for

30 min with rotation. Thereafter, the supernatant (unbound fraction) was removed. The beads were washed three times each with 3× SSC, 1× SSC, and 0.1× SSC for 2 min, rotating. Any remaining liquid was removed and beads were suspended in RNase-free water. RNA was eluted with TURBO DNase (Invitrogen) at room temperature for 45 min. For RNA purification, TRIzol Reagent (Invitrogen) and chloroform were added to all samples and the upper aqueous phase was further purified with RNA Clean & Concentrator kit (Zymo research) following the manufacturer's protocol.

mLeu-TTG ASO: /5Biosg/AAATGGGTGTCAGAAGTGGGATTCGAACCC

mAsp-GAY ASO: /5Biosg/AAATGGCTCCCCGTCGGGGAATCGAA

mGly-GGY ASO: /5Biosg/AAATGGTGCATGGGCCGGGAATCGAA

**LC–MS/MS analysis of nucleosides**

Untreated or chemically treated oligonucleotides (50–150 ng) were diluted in 17 μL nuclease-free water, denatured at 95 °C for 5 min, and immediately placed on ice for 2 min. Nuclease P1 (1 μL, 1 U/μL; Wako) and 2 μL of 100 mM ammonium acetate were added, and the samples were incubated at 42 °C for 2–4 h. Subsequently, 1 U FastAP thermosensitive alkaline phosphatase and 2.7 μL of 10× FastAP buffer (Thermo Fisher Scientific) were added, followed by incubation at 37 °C for 2–4 h. The digested samples were brought to a final volume of 60 μL with nuclease-free water and filtered through a 0.22 μm membrane filter. Nucleosides were analyzed using a Agilent 6460 Triple Quad LCMS system. For each analysis, 10 μL of the digested sample was injected and separated by reversed-phase ultra-high-performance liquid chromatography on a C18 column (Agilent). Nucleosides were detected in multiple-reaction monitoring mode using the precursor-to-product ion transitions. The channels of nucleosides were set as below:

2'-*O*-methyl-5-hydroxymethylcytidine (hm$^5$C*m*): 288.0→142.0

5-hydroxymethylcytidine (hm$^5$C): 274.0→142.0

2'-*O*-methyl-5-methylcytidine (m$^5$C*m*): 272.0→126.0

5-methylcytidine (m$^5$C): 258.0→126.0

2'-*O*-methylcytidine (C*m*): 258.2→112.1

cytidine (C): 244.0→112.0

2'-deoxy-5-hydroxymethylcytidine (5hmC): 258.1→142.1

2'-deoxy-5-hydroxymethyluridine (5hmU): 259.1→143.1

biotin-cysteamine-labeled 2'-deoxy-5-hydroxymethylcytidine (bsmC) 543.1→427.1

biotin-cysteamine-labeled 2'-deoxy-5-hydroxymethyluridine (bsmU): 544.1→428.1

**Preparation of DNA spike-ins**

All DNA templates and primers were purchased from IDT. The DNA 5hmC incorporated spike-in was synthesized by primer extension with ATP, GTP, 5hmCTP and TTP, using the template and primer as below.

DNA 5hmC template:

ACCCCTCACTACACCCCTCAACCATAGTCGCCACTGCGAGAAAGGTCCATCAGTACTGACAGTAGCATG
TACGTGACTCGCATATGCGATCGGATGGCACTACACATCTAGATCGGAAGAGCACAAACCTGAGTCACC
GTACCACACCTCTTAACTTCATTCAC

DNA 5hmC primer:

GTGAATGAAGTTAAGAGGTGTGGTACGGTGACTCAGGTTTGTGCTCTTCCGATCTAGATGTGTAGTGCCA
TCCGATCGCATATGCGAGTCACGTACATGCTACTGTCAGTACTGATGGACCTTTCTCGCAGTGGC

For the DNA C spike-in, the dsDNA oligonucleotides were purchased directly. Both dsDNA spike-ins were annealed, quantified using the Qubit dsDNA High Sensitivity (HS) Assay, and subsequently mixed at a 1:1 molar ratio.

DNA C sequence:

GTGAGTGGAGTTGAGAGGTGTGGTACGGTGACTCAGGTTTGTGCTCTTCCGATCTAGATGTGTAGTGCCATCCGATCGCATATGCGAGTCACGTACATGCTACTGTCAGTACTGATGGACCTTTCTCGCAGTGGCGACTATGGTTGAGGGGTGTAGTGAGGGGT

DNA 5hmC sequence:

GTGAATGAAGTTAAGAGGTGTGGTACGGTGACTCAGGTTTGTGCTCTTCCGATCTAGATGTGTAGTGCCATCCGATCGCATATGCGAGTCACGTACATGCTACTGTCAGTACTGATGGACCTTTCTCGCAGTGGCGA5hmCTATGGTTGAGGGGTGTAGTGAGGGGT

**Preparation of RNA spike-ins**

Both RNA spike-ins were synthesized by in vitro transcription (IVT) using an IVT kit (New England Biolabs). For the 5hmC-containing RNA spike-in, approximately 10% 5hmC incorporation was achieved by supplying $hm^5CTP$ (APExBIO) and CTP at a 1:9 molar ratio during IVT.

RNA C template:

ACCCCTCACTACACCCCTCAACCATAGTCGCCACTGCGAGAAAGGTCCATCAGTACTGACAGTAGCATGTACGTGACTCGCATATGCGATCGGATGGCACTACACATCTAGATCGGAAGAGCACAAACCTGAGTCACCGTACCACACCTCTCAACTCCACTCACTATAGTGAGTCGTATTA

RNA $hm^5C$ template:

ACCCCTCACTACACCCCTCAACCCGCTGATACACTGCGAGAAAGGTCCATCAGTACTGACAGTAGCATGTACGTGACTCGCATATGCGATCGGATGGCACTACACATCTAGATCGGAAGAGCACAAACCTGAGTCACCGTATCTCCACACCCAACTCCACTCACTATAGTGAGTCGTATTA

RNA C sequence:

TAATACGACTCACTATAGTGAGTGGAGTTGAGAGGTGTGGTACGGTGACTCAGGTTTGTGCTCTTCCGATCTAGATGTGTAGTGCCATCCGATCGCATATGCGAGTCACGTACATGCTACTGTCAGTACTGATGGACCTTTCTCGCAGTGGCGACTATGGTTGAGGGGTGTAGTGAGGGGT

RNA $hm^5C$ sequence:

TAATACGACTCACTATAGTGAGTGGAGTTGGGTGTGGAGATACGGTGACTCAGGTTTGTGCTCTTCCGATCTAGATGTGTAGTGCCATCCGATCGCATATGCGAGTCACGTACATGCTACTGTCAGTACTGATGGACCTTTCTCGCAGTGTATCAGCGGGTTGAGGGGTGTAGTGAGGGGT

T7 primer: TAATACGACTCACTATA

RT primer: ACCCCTCACTACACCCCTCAAC

PCR primer RNA C F: GTGAGTGGAGTTGAGAGGTGTGG

PCR primer RNA C R: ACCCCTCAACCATAGTCGCC

PCR primer RNA $hm^5C$ F: GTGAGTGGAGTTGGGTGTGGAGA

PCR primer RNA $hm^5C$ R: ACCCCTCAACCCGCTGATAC

**RNA $hm^5C$ immunoprecipitation (vs BALT)**

Condition 1: hMeRIP[2]

100 ng of denatured RNA spike-ins (hm$^5$C-containing: non-hm$^5$C-containing at 1:100 molar ratio) were subjected to immunoprecipitation with anti-5hmC (Active Motif, 39769) with an equal volume of 2× immunoprecipitation buffer was added (40 mM Tris pH 7.4, 1 mM EDTA pH 8.0, 700 mM NaCl, 0.2% NP-40) at 4 ˚C overnight. Immunoprecipitated RNA was recovered using protein G Dynabeads. Immunoprecipitated RNA was washed three times with wash buffer (20 mM Tris-HCl pH 7.4RT, 0.5 mM EDTA, 350 mM NaCl, 0.1% NP-40).

Condition 2: BALT and streptavidin enrichment

100 ng of denatured RNA spike-ins (hm$^5$C-containing: non-hm$^5$C-containing at 1:100 molar ratio) were subjected to general BALT treatment protocol in 20 μL scale. After the treatment, ssDNA was desalted using a Bio-Spin P-6 Gel Column (Bio-Rad) and purified by Oligo Clean & Concentrator (Zymo Research). Biotin-labeled RNA was enriched using Dynabeads MyOne Streptavidin C1 (Invitrogen). Streptavidin beads (5 μL) were washed and blocked with RNase-free BSA for 30 min at room temperature. Treated RNA was incubated with the blocked beads in binding buffer for 30 min at room temperature. Beads were subsequently subjected to two times of high-salt wash at pH 7.5 (5 mM Tris pH 7.5, 0.5 mM EDTA, 1 M NaCl and 0.2% Tween 20) and two times of high-salt wash at pH 7.5 without NaCl (5 mM Tris pH 7.5, 0.5 mM EDTA and 0.2% Tween 20) to remove nonspecifically bound RNA. After the final wash, bead-bound RNA was resuspended in 10 μL nuclease-free water and heated at 95 °C for 10 min. The beads were immediately removed on a magnetic rack, and the RNA-containing supernatant was transferred to a new RNase-free tube.

Enriched RNA was purified by TRIzol reagent (Invitrogen), added with 200 μL chloroform, vigorously vortexed and centrifuged at 16,000× g for 1 min and the supernatant was mixed with one volume of 100% ethanol. RNA was bound to the column and washed with washing buffer and eluted with 50 μl RNase-free water, followed by RT–qPCR with specific RT primer. Reverse transcription was performed by Superscript IV kit (Invitrogen).

**RT–qPCR**

RNA was reverse transcribed using PrimeScript RT Master Mix (Takara Bio) with both oligo(dT) and random hexamer primers according to the manufacturer's instructions. The resulting cDNA was analyzed by quantitative PCR (qPCR) on a LightCycler 96 system (Roche) using FastStart Essential DNA Green Master (Roche) and gene-specific primers. Relative enrichment fold or recovery were determined using the ΔΔCt method. All PCR primers were purchased from IDT. [3,4]

L1Md_A_F: GGATTCCACACGTGATCCTAA

L1Md_A_R: TCCTCTATGAGCAGACCTGGA

L1Md_Gf_F: CTCCTTGGCTCCGGGACT

L1Md_Gf_R: CAGGAAGGTGGCCGGTTGT

L1Md_Tf_F: CAGCGGTCGCCATCTTG

L1Md_Tf_R: CACCCTCTCACCTGTTCAGACTAA

IAPEz_F: CTCCATGTGCTCTGCCTTCC

IAPEz_R: CCCCGTCCCTTTTTTAGGAGA

**mESCs culture and isolation of genomic DNA and total RNA**

The mESCs cell line ES-E14TG2a was purchased from ATCC and grown on gelatincoated plates in complete DMEM (Invitrogen Cat. No. 11995) supplemented with 15% vol/vol FBS (Gibco), 1% penicillin/streptomycin (Gibco), 1.25× nucleoside (MilliporeSigma), 62.5 mM 2-mercaptoethanol (Thermo Fisher Scientific), 1.25× nonessential amino acids (Gibco), 104 U/mL leukemia inhibitory factor (LIF) (MilliporeSigma), 0.289 mg per 500 mL of PD0325901 (STEMCELL Technologies), 0.83 mg per 500 mL of CHIR99021 (STEMCELL Technologies), and 2.5 mg/L Plasmocin

Prophylactic (Invitrogen). The cells were cultured at 37 °C with 5.0% $CO_2$ and passaged every 2 days. Authentication and mycoplasma contamination testing were performed within the lab to confirm cell identity and purity. After the cells were harvested via centrifugation for 3 min at 1000×g, gDNA was extracted with a PureLink Genomic DNA Mini Kit (Invitrogen) following the manufacturer's protocol. total RNA was isolated from the cells using TRIzol reagent (Ambion by Life Technologies) and Direct-zol RNA miniprep kit (Zymo Research) following the manufacturer's protocol. In brief, cells from 10-cm plates were suspended in 1 ml of TRIzol reagent (Invitrogen), added with 200 µL chloroform, vigorously vortexed and centrifuged at 16,000× g for 1 min and the supernatant was mixed with one volume of 100% ethanol. RNA was bound to the column and treated with DNase I at room temperature for 15 min. RNA was washed with washing buffer and eluted with 50 µl RNase-free water.

**Isolation of cell-free DNA**

Cell-free DNA (cfDNA) was extracted and purified from pooled human plasma (Innovative Research Inc., Cat. #IPLAWBK2E) using the Qiagen QIAamp Circulating Nucleic Acid kit followed by Ampure beads size selection to remove gDNA contamination. For library preparation, directly perform end repair and A-tailing without fragmentation.

**BALT-seq (vs 5hmC-Seal) DNA library preparation**

Mouse Genomic DNA (0.1–1 µg) was fragmented using NEBNext dsDNA Fragmentase (New England Biolabs) in a 20 µL reaction containing 2 µL Fragmentase Reaction Buffer v2 and 2 µL dsDNA Fragmentase. The reaction was incubated at 37 °C for 30 min and quenched by addition of 5 µL of 0.5 M EDTA (pH 8.0). Fragmented DNA was purified using a DNA Clean & Concentrator-5 kit (Zymo Research) and eluted in nuclease-free water. 10 ng fragmented DNA was used for next step, quantified using the Qubit dsDNA High Sensitivity (HS) Assay.

DNA end repair and adaptor ligation were performed using the NEBNext Ultra II DNA Library Prep Kit for Illumina (New England Biolabs). Briefly, 5–100 ng of fragmented DNA was subjected to end repair at 20 °C for 30 min followed by 65 °C for 30 min. The NEBNext adaptor was diluted 10-fold to 1.5 µM and ligated to the repaired DNA at 20 °C for 15 min, followed by USER enzyme treatment at 37 °C for 15 min. Adaptor-ligated DNA was purified using 0.9× AMPure XP beads (Beckman Coulter) and eluted in 25 µL nuclease-free water.

Condition 1: BALT

Fresh 2× sodium bisulfite solution (2× SBS) was prepared by dissolving 190 mg sodium metabisulfite in 980 µL nuclease-free water followed by addition of 20 µL of 37% HCl. Adaptor-ligated DNA was denatured at 95 °C for 5 min and immediately placed on ice. Biotin-cysteamine (2 mg) was dissolved in 25 µL freshly prepared 2× SBS and preheated at 80 °C for 5 min. Denatured DNA (25 µL) was combined with 25 µL of the biotin-cysteamine/2× SBS solution and incubated at 40 °C for 30 min.

The reaction was desalted using a Bio-Spin P-6 Gel Column (Bio-Rad). The recovered DNA was subsequently desulfonated using the EZ DNA Methylation-Gold Kit (Zymo Research). Samples were incubated with M-Desulphonation Buffer at room temperature for 20 min, washed extensively, and eluted in 30 µL nuclease-free water. An aliquot (1 µL) was retained as the input control before affinity enrichment.

Condition 2: 5hmC-Seal

Adaptor-ligated DNA (20 µL) was supplemented with 2.5 µL T4 phage $\beta$-glucosyltransferase (T4 $\beta$GT; Thermo Fisher Scientific), 2.5 µL Epi-buffer, and 1 µL of 3 mM UDP-azide-glucose (Active Motif) in a final reaction volume of 26 µL. The reaction mixture was incubated at 37 °C for 2 h. The DNA was subsequently purified using a DNA Clean & Concentrator-5 kit (Zymo Research) according to the manufacturer's instructions and eluted in 30 µL nuclease-free water preheated to 56 °C. For biotin conjugation, 1 µL of 4.5 mM DBCO-biotin in DMSO was added to the purified DNA, and the reaction mixture was incubated at 37 °C for 2 h. The biotinylated DNA was purified again using a DNA Clean & Concentrator-5 kit and eluted in 30 µL nuclease-free water. An aliquot (1 µL) was retained as the input control before affinity enrichment.

Biotin-labeled DNA was enriched using Dynabeads MyOne Streptavidin C1 (Invitrogen). Streptavidin beads (2.5 µL per sample) were washed and blocked with salmon sperm DNA for 30 min at room temperature. The treated DNA was incubated with the blocked beads in binding buffer at room temperature for 30 min. Beads were subsequently subjected to two times of high-salt wash at pH 7.5 (5 mM Tris pH 7.5, 0.5 mM EDTA, 1 M NaCl and 0.2% Tween 20), two times of high-salt wash at pH 7.5 without NaCl (5 mM Tris pH 7.5, 0.5 mM EDTA and 0.2% Tween 20), two times of low-salt wash at pH 9.0 (5 mM Tris pH 9.0, 0.5 mM EDTA, 1 M NaCl and 0.2% Tween 20) and two times of low-salt wash (5 mM Tris pH 9.0, 0.5 mM EDTA and 0.2% Tween 20) to remove nonspecifically bound DNA. After the final wash, the beads were resuspended in 10 µL nuclease-free water.

Sequencing libraries were amplified directly from the bead-bound DNA using NEBNext Ultra II Q5 Master Mix with the corresponding i5 and i7 indexing primers. The optimal number of amplification cycles was determined by qPCR; approximately 16 cycles were typically used for libraries generated from ~10 ng input DNA. PCR products were purified using 0.9× AMPure XP beads and eluted in 20 µL nuclease-free water.

**BALT-seq RNA library preparation**

Total RNA (~5 µg) was fragmented in a 20 µL reaction containing 10 mM Tris-HCl (pH 7.5) and 10 mM $ZnCl_2$. RNA was incubated at 70 °C for 5 min, and fragmentation was immediately quenched by addition of 5 µL of 500 mM EDTA (pH 8.0). Fragmented RNA was purified using the RNA Clean & Concentrator-5 kit (Zymo Research) and eluted in 25 µL nuclease-free water. The indicated RNA spike-in was added after purification.

Fresh 2× SBS was prepared by dissolving 190 mg sodium metabisulfite in 980 µL nuclease-free water followed by addition of 20 µL of 37% HCl. Biotin-cysteamine (2 mg) was dissolved in 100 µL freshly prepared 2× SBS and preheated at 70 °C for 10 min. Fragmented RNA was separately heated at 70 °C for 3 min and immediately combined with an equal volume of the preheated biotin-cysteamine/2× SBS solution. The 50 µL reaction was incubated at 40 °C for 30 min.

The labeling reaction was desalted using a Bio-Spin P-6 Gel Column (Bio-Rad). The recovered RNA was purified using an RNA Clean & Concentrator-5 column and treated with RNA Desulphonation Buffer at room temperature for 40 min. Following desulfonation and extensive washing, RNA was eluted in 21 µL nuclease-free water. An aliquot (1 µL) was retained as the input control before affinity enrichment.

Biotin-labeled RNA was enriched using Dynabeads MyOne Streptavidin C1 (Invitrogen). Streptavidin beads (5 µL per sample) were washed and blocked with RNase-free BSA for 30 min at room temperature. Treated RNA was incubated with the blocked beads in binding buffer for 30 min at room temperature. The beads were then subjected to the same sequential high- and low-salt washing buffer at pH 7.5 and pH 9.0 used for DNA BALT-seq. After the final wash, bead-bound RNA was resuspended in 9 µL nuclease-free water and heated at 95 °C for 10 min. The beads were immediately removed on a magnetic rack, and the RNA-containing supernatant was transferred to a new RNase-free tube.

RNA sequencing libraries were prepared using the Takara library construction workflow Option 2 (Takara Bio) without an additional fragmentation step, as the RNA had been fragmented before BALT treatment.

**General Synthesis Remarks**

All purchased solvents and substrates were used without further treatment unless otherwise stated. Yields refer to chromatographically and spectroscopically ($^1$H NMR) homogeneous material, unless otherwise stated. Chemical reactions were monitored by thin layer chromatography (TLC). Analytical thin-layer chromatography (TLC) was conducted using 0.2 mm commercial silica gel plates (silica gel 60, $F_{254}$, EMD chemical). Analytical reversed-phase thinlayer chromatography (TLC) was carried out using 0.2 mm commercial C18 silica gel plates (silica gel 60 RP-18, $F_{254}$S). Vials (15 Å~ 45 mm 1 dram (4 mL) /17 Å~ 60 mm 3 dram (7.5 mL) with PTFE lined cap attached) were purchased from Qorpak and used as received. Nuclear magnetic resonance spectra ($^1$H NMR and $^{13}$C NMR) were recorded with Bruker Model DMX 600 (600 MHz, $^1$H at 600 MHz, $^{13}$C at 151 MHz) or 400 (400 MHz, $^1$H at 400 MHz, $^{13}$C at 101 MHz). Unless otherwise noted, all spectra were acquired in $d^6$-DMSO. Chemical shifts are reported in parts per million (ppm, δ) and are referenced to the residual solvent (DMSO, $\delta$ = 2.50 ppm ($^1$H) and 39.52 ppm ($^{13}$C)). Coupling constants were reported in Hertz (Hz). Data for $^1$H NMR spectra were reported as follows: chemical shift (ppm, referenced to protium, s = singlet, d = doublet, t = triplet, q = quartet, quin = quintet, oct = octet, dd = doublet of doublets, td = triplet of doublets, ddd = doublet of doublet of doublets, m = multiplet, coupling constant (Hz), and integration). Column chromatography was performed using E. Merck silica (60, particle size 0.043–0.063 mm), and pTLC was performed on Merck silica plates (60F-254). High-resolution mass spectra (HRMS) were recorded on an Agilent 6530 LC Q-TOF mass spectrometer using electrospray ionization with fragmentation voltage set at 75-130 V and processed with an Agilent MassHunter Operating System.

**Genreral procedure for synthesis of biotin-thiol** [5]

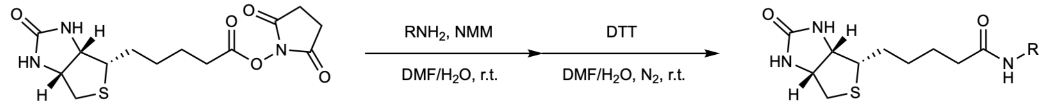


To a 100 mL flask, mercaptamine (1.0 mmol) and NMM (1.1 mmol) was added in DMF/$H_2O$ (5:1, 30 mL). After mercaptamine was dissolved, NHS-biotin was then added, and the reaction mixture was stirred for 8 h at room temperature. White precipitation occurred, which is the corresponding disulfide oxidized from the desired product. To reduce the disulfide, dithiothreitol (2.0 mmol) was added and then the reaction mixture was kept under an nitrogen atmosphere for 2 h until the mixture was transparent. The solvent was evaporated under vacuum, and the residue was dissolved in MeOH. Silica gel was added, and then MeOH was evaporated under vacuum so that the mixture can be purified via column chromatography (DCM/MeOH, gradient from 100:1 to 10:1) to provide the desired product as white solid. Ellman's reagent (5 mg dissolved in DCM/MeOH) can be used to detect thiol groups on TLC plate.

**Synthesis of biotin-cysteamine**

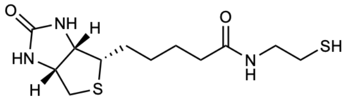

*N*-(2-mercaptoethyl)-5-((3a*S*,4*S*,6a*R*)-2-oxohexahydro-1*H*-thieno[3,4-*d*]imidazol-4-yl)pentanamide

According to general procedure, NHS-biotin (341 mg, 1.0 mmol), cysteamine hydrochloride (114 mg, 1.0 mmol) and NMM (222 mg, 2.2 mmol) was added in the first step, and then dithiothreitol (308 mg, 2.0 mmol) was added. After steps of treatments as described, the mixture was purified by column chromatography to yield the compound as white solid (246 mg, 81%).

$^1$H NMR (600 MHz, DMSO) δ 7.95 (t, J = 5.7 Hz, 1H), 6.42 (s, 1H), 6.36 (s, 1H), 4.31 (dd, J = 7.7, 5.0 Hz, 1H), 4.14 (ddd, J = 7.0, 4.5, 1.9 Hz, 1H), 3.19 (q, J = 6.4 Hz, 2H), 3.11 (ddd, J = 8.8, 6.1, 4.3 Hz, 1H), 2.83 (dd, J = 12.4, 5.1 Hz, 1H), 2.58 (d, J = 12.5 Hz, 1H), 2.55 – 2.48 (m, 2H), 2.34 (t, J = 8.0 Hz, 1H), 2.07 (t, J = 7.4 Hz, 2H), 1.66 – 1.43 (m, 4H), 1.38 – 1.25 (m, 2H).

$^{13}$C NMR (151 MHz, DMSO) δ 172.57, 163.18, 61.51, 59.67, 55.88, 42.51, 40.32, 35.61, 28.66, 28.50, 25.71, 24.01.

HRMS (ESI-TOF) *m/z* calcd for $C_{12}H_{21}N_3O_2S_2$ 303.1075; found 303.1077.

**Synthesis of biotin-PEG-SH**

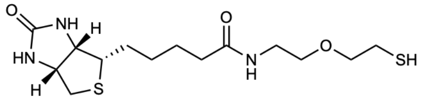

*N*-(2-(2-mercaptoethoxy)ethyl)-5-((3a*S*,4*S*,6a*R*)-2-oxohexahydro-1*H*-thieno[3,4-*d*]imidazol-4-yl)pentanamide

According to general procedure, NHS-biotin (170 mg, 0.5 mmol), 2-(2-aminoethoxy)ethane-1-thiol hydrochloride (79 mg, 0.5 mmol) and NMM (111 mg, 1.1 mmol) was added in the first step, and then dithiothreitol (153 mg, 1.0 mmol) was added. After steps of treatments as described, the mixture was purified by column chromatography to yield the compound as white solid (125 mg, 72%).

$^{1}H$ NMR (600 MHz, DMSO) δ 7.87 (t, *J* = 5.7 Hz, 1H), 6.43 (d, *J* = 2.5 Hz, 1H), 6.38 (s, 1H), 4.31 (dd, *J* = 7.8, 5.0 Hz, 1H), 4.13 (ddd, *J* = 7.0, 4.4, 2.1 Hz, 1H), 3.49 (t, *J* = 6.6 Hz, 2H), 3.40 (t, *J* = 5.9 Hz, 2H), 3.19 (q, *J* = 5.7 Hz, 2H), 3.10 (ddd, *J* = 8.6, 6.1, 4.4 Hz, 1H), 2.82 (dd, *J* = 12.5, 5.1 Hz, 1H), 2.63 – 2.59 (m, 3H), 2.33 (t, *J* = 8.0 Hz, 1H), 2.07 (t, *J* = 7.4 Hz, 2H), 1.65 – 1.45 (m, 3H), 1.49 – 1.20 (m, 3H).

$^{13}C$ NMR (151 MHz, DMSO) δ 173.24, 163.22, 72.39, 69.25, 69.24, 61.52, 59.68, 55.90, 40.33, 38.90, 35.58, 28.66, 28.51, 23.90.

HRMS (ESI-TOF) *m/z* calcd for $C_{14}H_{25}N_3O_3S_2$ 347.1337; found 347.1336.

**Synthesis of biotin-cysteine**

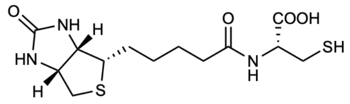


(5-((3a*S*,4*S*,6a*R*)-2-oxohexahydro-1*H*-thieno[3,4-*d*]imidazol-4-yl)pentanoyl)-*L*-cysteine

According to general procedure, NHS-biotin (341 mg, 1.0 mmol), *L*-cysteine (121 mg, 1.0 mmol) and NMM (111 mg, 1.1 mmol) was added in the first step, and then dithiothreitol (308 mg, 2.0 mmol) was added. After the solvent was evaporated, impurity in the residual was dissolved in MeOH by ultrasound treatment to yield the compound as white solid (282 mg, 81%).

$^{1}H$ NMR (600 MHz, DMSO) δ 12.80 (s, 1H), 8.12 (d, J = 8.0 Hz, 1H), 6.41 (s, 1H), 6.36 (s, 1H), 4.38 (td, J = 7.7, 4.6 Hz, 1H), 4.31 (dd, J = 7.7, 5.1 Hz, 1H), 4.17 – 4.11 (m, 1H), 3.14 – 3.07 (m, 1H), 2.89 – 2.70 (m, 3H), 2.58 (d, J = 12.5 Hz, 1H), 2.41 (t, J = 8.7 Hz, 1H), 2.17 (t, J = 7.4 Hz, 2H), 1.67 – 1.43 (m, 4H), 1.41 – 1.27 (m, 2H).

$^{13}C$ NMR (151 MHz, DMSO) δ 172.77, 172.27, 163.18, 61.51, 59.67, 55.90, 54.77, 40.33, 35.26, 28.56, 28.47, 26.05, 25.66.

HRMS (ESI-TOF) *m/z* calcd for $C_{13}H_{21}N_3O_4S_2$ 347.0973; found 347.0974.

**Synthesis of 2'-*O*-methyl-5-hydroxymethylcytidine** [6]

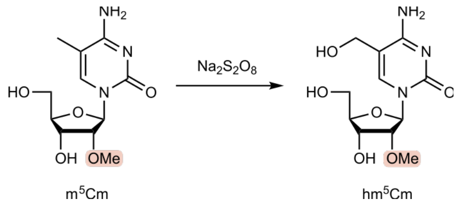


2'-*O*-methyl-5-methylcytidine was purchased from BIOSYNTH. In brief, 2'-*O*-methyl-5-methylcytidine (45 mg, 0.16 mmol) was suspended in DBPS and sodium persulfate (40 mg, 0.16 mmol) was added and the reaction mixture was heated at 80 °C for 4 h. The reaction mixture was concentrated under reduced pressure and the crude product was purified by silica gel plates (20 % MeOH in DCM) to yield the product 2'-*O*-methyl-5-hydroxymethylcytidine as white solid (8 mg, 17 %).

$^{1}H$ NMR (400 MHz, DMSO) δ 7.88 (s, 1H), 7.37 (s, 1H), 6.71 (s, 1H), 5.87 (d, *J* = 4.2 Hz, 1H), 5.18 – 5.05 (m, 3H), 4.17 (d, *J* = 5.4 Hz, 2H), 4.08 (q, *J* = 5.6 Hz, 1H), 3.82 (dt, *J* = 5.9, 3.1 Hz, 1H), 3.72 – 3.62 (m, 2H), 3.56 (ddd, *J* = 12.1, 5.1, 3.3 Hz, 1H), 3.38 (s, 3H).

**Data Analysis of DNA BALT-seq and 5hmC-Seal-seq**

Paired-end sequencing reads were adapter- and quality-trimmed with Trim Galore. Trimmed reads were aligned to the mouse mm10 genome using Bowtie2 (v2.5.4) in end-to-end mode with sensitive settings (--end-to-end --sensitive -k 5 -X 1000). Secondary alignments were removed using SAMtools (samtools view -F 256) to retain uniquely mapped reads.

PCR duplicates were removed from uniquely mapped BAM files using Picard MarkDuplicates with REMOVE_DUPLICATES=true. The duplicate-removed BAMs were further filtered to remove mitochondrial and ENCODE blacklist-region reads. 5hmC-enriched regions were identified using MACS2 broad peak calling against the matched input control BAM (--broad --broad-cutoff 0.05 –nomodel –nolambda -g mm -B --keep-dup all).

Single-base-resolution 5hmC sites detected by TAB-seq and oxBS-seq were downloaded from GEO datasets GSM882244 and GSM4194641 and lifted over from mm9 to mm10. BALT-seq read coverage around these 5hmC sites was calculated using deepTools.

**Data Analysis of RNA BALT-RNA data**

Paired-end sequencing reads were adapter- and quality-trimmed with Trim Galore. Trimmed reads were aligned to the mouse reference genome mm10 using HISAT2 (v2.2.1, --no-softclip -k 10) with known splice sites from GENCODE release M25. Secondary alignments were removed with SAMtools. PCR duplicates were removed using Picard MarkDuplicates with REMOVE_DUPLICATES=true. Reads overlapping ENCODE blacklist regions from the mm10 reference index were removed with BEDTools.

For peak discovery, IP and matched input BAM files were normalized by downsampling to approximately 2.9 million and 4.0 million reads, respectively. Exonic enrichment peaks were identified using ExonPeak with GENCODE M25 gene annotation. For non-exonic regions, reads were first restricted to GENCODE-defined non-exonic intervals and separated by strand using paired-end SAM flags. Strand-specific non-exonic peaks were then called with MACS2 using matched input controls and the following parameters: -f BAM -g mm --nomodel --nolambda -B -q 0.01 --keep-dup all --extsize 150. Plus- and minus-strand peak calls were merged into strand-aware peak sets, which were further annotated against the mm10 genome. Read counts over peaks and RepeatMasker repeat annotations were quantified separately using featureCounts v2.0.3.

### NMR Spectra

#### $^{1}$H NMR of biotin-cysteamine

$^{1}$H NMR (600 MHz, DMSO) δ 7.95 (t, $J$ = 5.7 Hz, 1H), 6.42 (s, 1H), 6.36 (s, 1H), 4.31 (dd, $J$ = 7.7, 5.0 Hz, 1H), 4.14 (ddd, $J$ = 7.0, 4.5, 1.9 Hz, 1H), 3.19 (q, $J$ = 6.4 Hz, 2H), 3.11 (ddd, $J$ = 8.8, 6.1, 4.3 Hz, 1H), 2.83 (dd, $J$ = 12.4, 5.1 Hz, 1H), 2.58 (d, $J$ = 12.5 Hz, 1H), 2.55 – 2.48 (m, 2H), 2.34 (t, $J$ = 8.0 Hz, 1H), 2.07 (t, $J$ = 7.4 Hz, 2H), 1.66 – 1.43 (m, 4H), 1.38 – 1.25 (m, 2H).

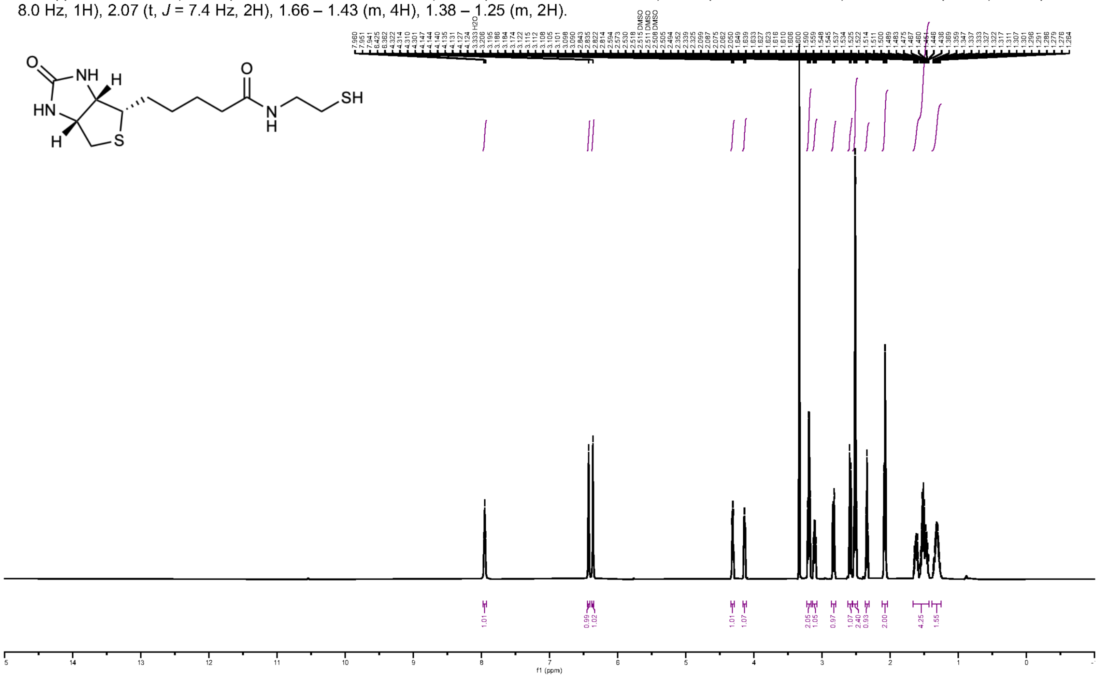


#### $^{13}$C NMR of biotin-cysteamine

$^{13}$C NMR (151 MHz, DMSO) δ 172.57, 163.18, 61.51, 59.67, 55.88, 42.51, 40.32, 35.61, 28.66, 28.50, 25.71, 24.01.

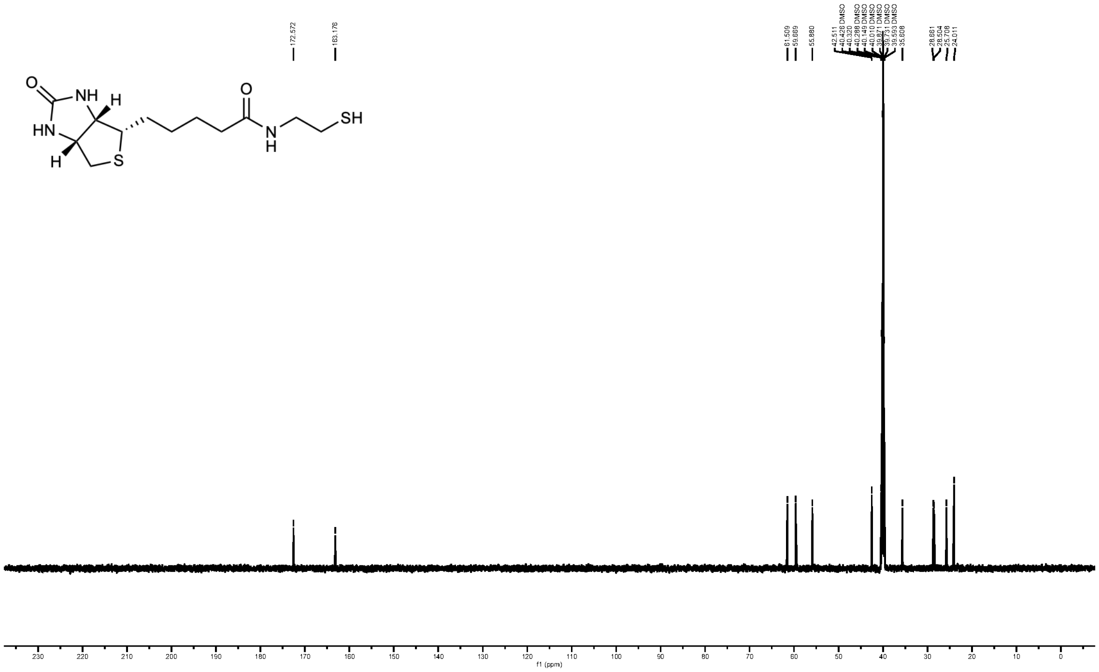

### $^{1}$H NMR of biotin-PEG-SH

$^{1}$H NMR (600 MHz, DMSO) δ 7.87 (t, *J* = 5.7 Hz, 1H), 6.43 (d, *J* = 2.5 Hz, 1H), 6.38 (s, 1H), 4.31 (dd, *J* = 7.8, 5.0 Hz, 1H), 4.13 (ddd, *J* = 7.0, 4.4, 2.1 Hz, 1H), 3.49 (t, *J* = 6.6 Hz, 2H), 3.40 (t, *J* = 5.9 Hz, 2H), 3.19 (q, *J* = 5.7 Hz, 2H), 3.10 (ddd, *J* = 8.6, 6.1, 4.4 Hz, 1H), 2.82 (dd, *J* = 12.5, 5.1 Hz, 1H), 2.63 – 2.59 (m, 3H), 2.33 (t, *J* = 8.0 Hz, 1H), 2.07 (t, *J* = 7.4 Hz, 2H), 1.65 – 1.45 (m, 3H), 1.49 – 1.20 (m, 3H).

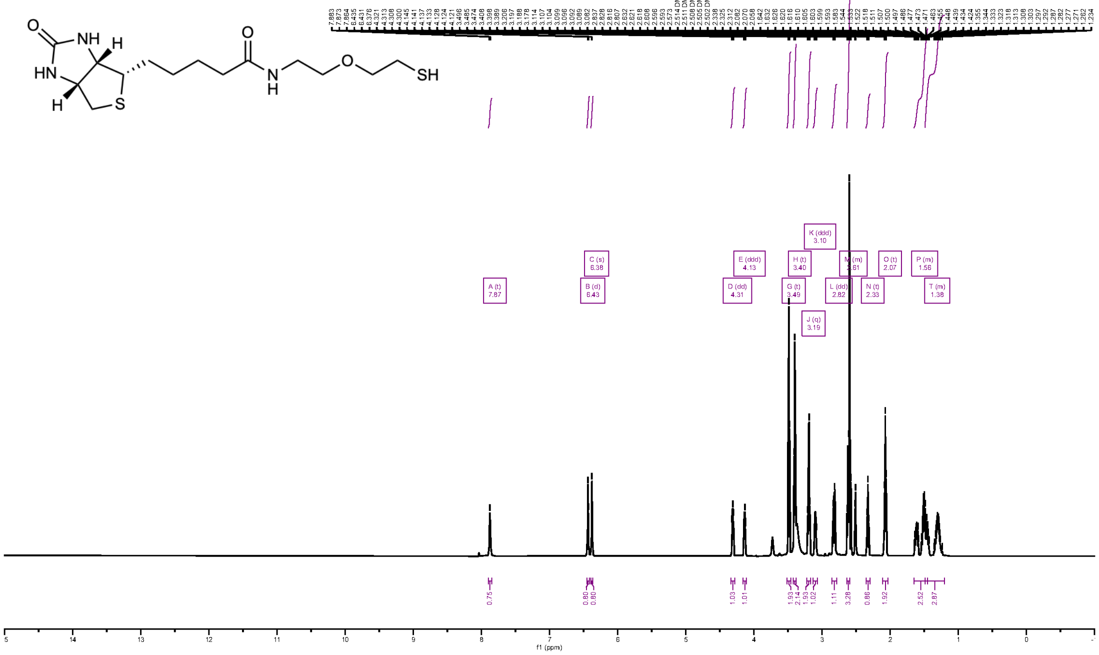

### $^{13}$C NMR of biotin-PEG-SH

$^{13}$C NMR (151 MHz, DMSO) δ 173.24, 163.22, 72.39, 69.25, 69.24, 61.52, 59.68, 55.90, 40.33, 38.90, 35.58, 28.66, 28.51, 23.90.

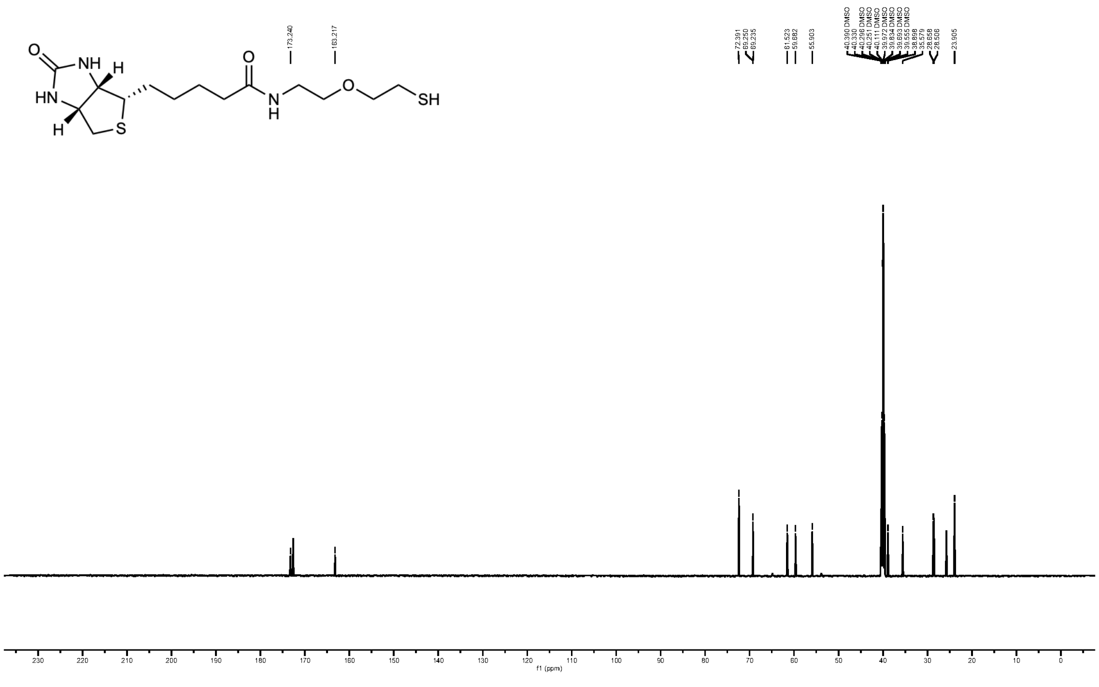

### [1]H NMR of biotin-cysteine

[1]H NMR (600 MHz, DMSO) δ 12.80 (s, 1H), 8.12 (d, *J* = 8.0 Hz, 1H), 6.41 (s, 1H), 6.36 (s, 1H), 4.38 (td, *J* = 7.7, 4.6 Hz, 1H), 4.31 (dd, *J* = 7.7, 5.1 Hz, 1H), 4.17 – 4.11 (m, 1H), 3.14 – 3.07 (m, 1H), 2.89 – 2.70 (m, 3H), 2.58 (d, *J* = 12.5 Hz, 1H), 2.41 (t, *J* = 8.7 Hz, 1H), 2.17 (t, *J* = 7.4 Hz, 2H), 1.67 – 1.43 (m, 4H), 1.41 – 1.27 (m, 2H).

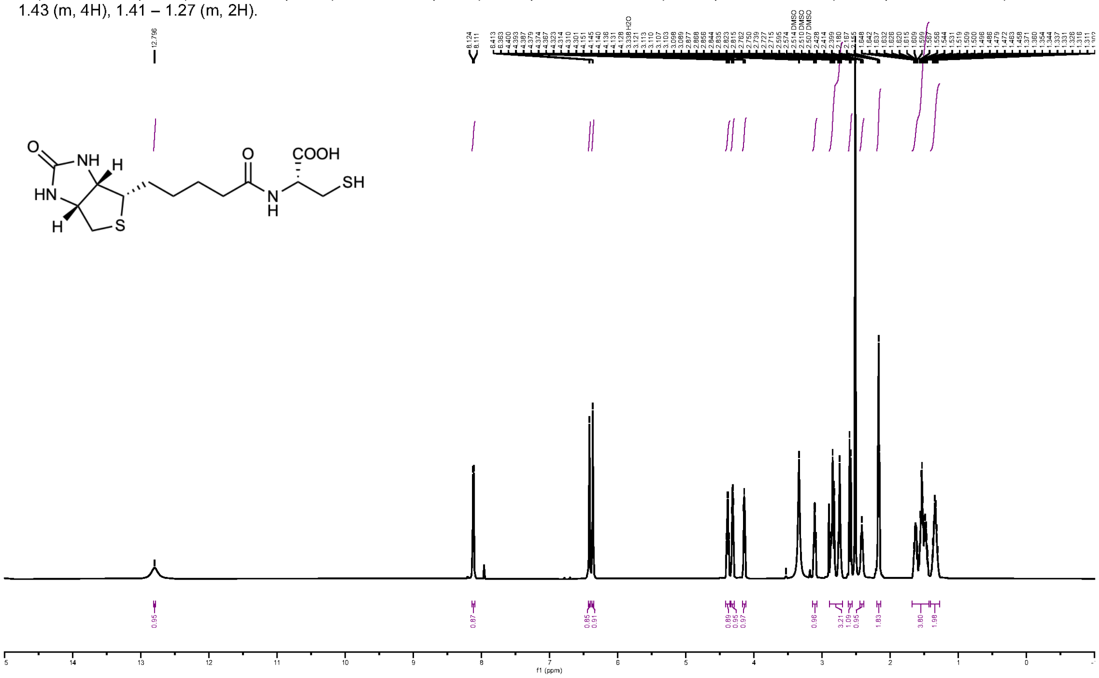


### [13]C NMR of biotin-cysteine

[13]C NMR (151 MHz, DMSO) δ 172.77, 172.27, 163.18, 61.51, 59.67, 55.90, 54.77, 40.33, 35.26, 28.56, 28.47, 26.05, 25.66.

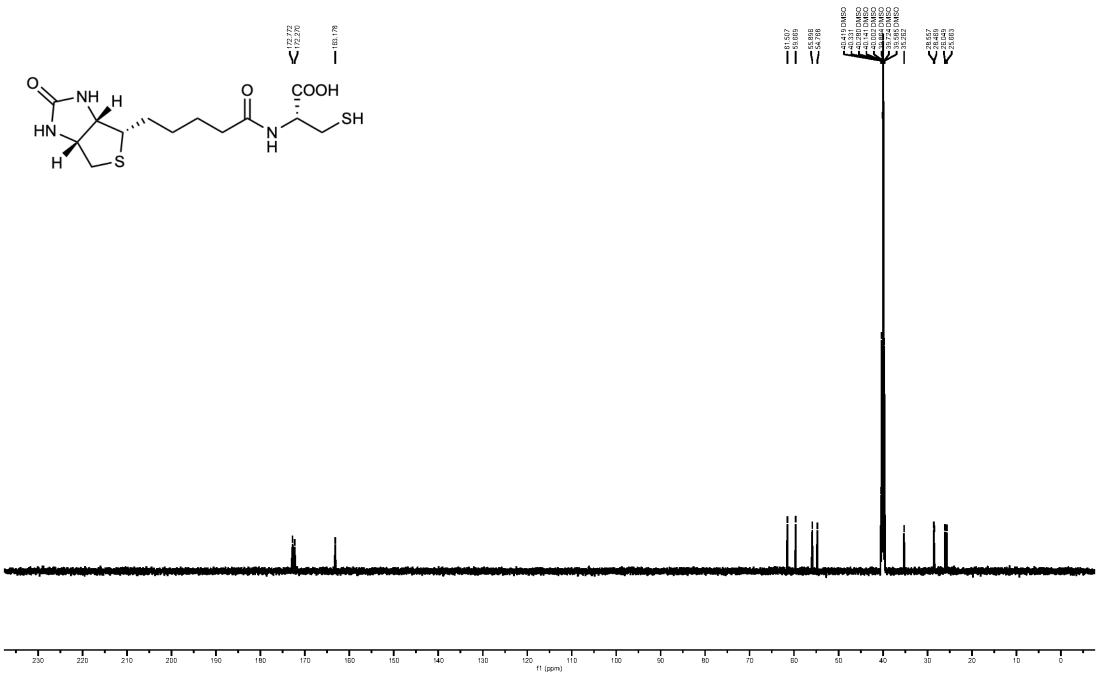

### $^{1}$H NMR of 2'-*O*-methyl-5-hydroxymethylcytidine

$^{1}$H NMR (400 MHz, DMSO) δ 7.88 (s, 1H), 7.37 (s, 1H), 6.71 (s, 1H), 5.87 (d, $J$ = 4.2 Hz, 1H), 5.18 – 5.05 (m, 3H), 4.17 (d, $J$ = 5.4 Hz, 2H), 4.08 (q, $J$ = 5.6 Hz, 1H), 3.82 (dt, $J$ = 5.9, 3.1 Hz, 1H), 3.72 – 3.62 (m, 2H), 3.56 (ddd, $J$ = 12.1, 5.1, 3.3 Hz, 1H), 3.38 (s, 3H).

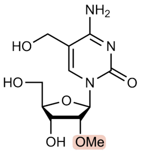

hm$^5$Cm

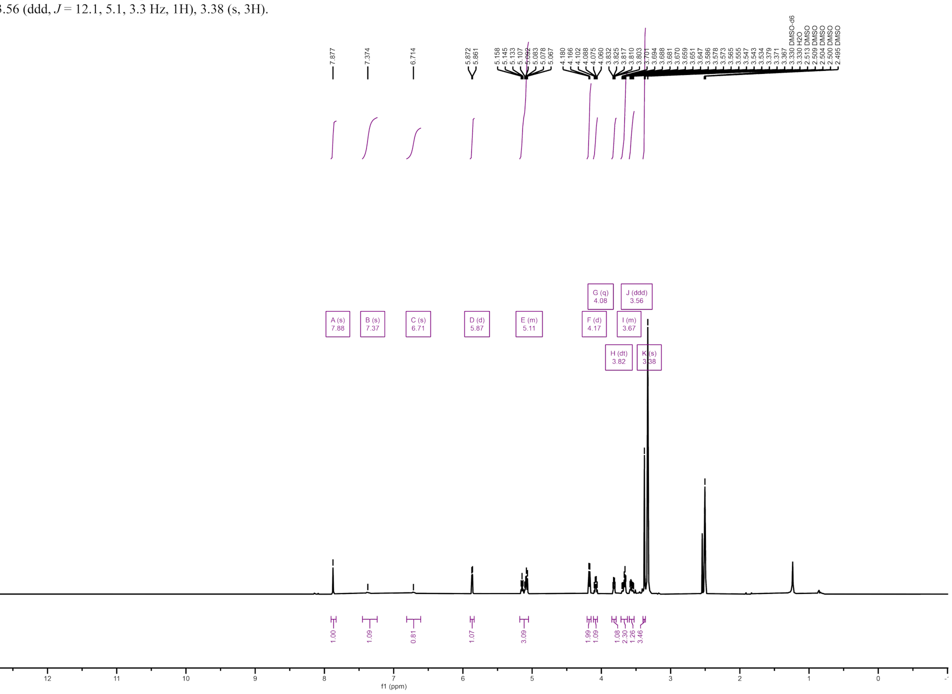
